# Plasticity of extrachromosomal DNA segregation during drug adaptation

**DOI:** 10.64898/2025.12.09.693337

**Authors:** Chikako Shibata, Kenichi Miyata, Kohei Kumegawa, Liying Yang, Ryu-Suke Nozawa, Reo Maruyama

## Abstract

Uneven segregation during mitosis is a striking feature of extrachromosomal DNA (ecDNA). Because ecDNA lacks a centromere, it is thought to segregate stochastically and randomly during cell division, thereby generating extensive intratumoral heterogeneity in genomic copy number. Several studies have reported that ecDNA copy numbers can readily change in response to drug treatment, which enables the cells to acquire drug resistance. However, the mechanisms underlying these dynamic changes remain poorly understood —particularly whether such copy-number changes result from static selection of pre-existing clones or from active reconfiguration under drug-induced stress. This key question remains unresolved, mainly due to the absence of technologies capable of tracking ecDNA copy number simultaneously across clones. To overcome this limitation, we developed a high-throughput framework that combines single-cell DNA sequencing with cellular barcoding for clonal tracking of ecDNA copy-number dynamics.

The results of single-cell cloning experiments revealed that not all clones exhibit identical segregation modes even under drug-free conditions. Clonal tracking under drug treatment showed that resistant populations do not simply emerge from pre-existing clones with favorable ecDNA states; instead, some clones seemed capable of actively reconfiguring their segregation behavior, possibly involving neuron-like alternation in microtubule organization and intracellular transport pathways, to generate drug-resistant cells. These findings suggest that, although ecDNA might segregate stochastically, it may undergo nonrandom, actively regulated segregation under drug stress. This discovery allows the therapeutically targeting of ecDNA segregation mechanisms to counteract adaptive drug resistance.

## Introduction

Cancer cells continually reshape their genomes to adapt and survive under stress. Among the most dynamic forms of genome organization is extrachromosomal DNA (ecDNA) —a circular, acentromeric DNA element, frequently carrying oncogenes^1,2^. A hallmark of ecDNA is its uneven segregation during mitosis, which generates striking heterogeneity in oncogene copy number within the tumor^3,4–7^. This heterogeneity has been implicated in drug resistance and rapid clonal evolution^4,8–11^.

Several studies have reported that, after drug exposure, resistant populations often show altered ecDNA copy numbers compared with the parental cells, suggesting that ecDNA dynamics contribute to adaptation^4,8–10^. However, it is unclear whether these changes only reflect the selection of pre-existing clones with favorable ecDNA states or arise from active reconfiguration of ecDNA content during stress. This question lies at the heart of understanding whether ecDNA segregation is purely stochastic or can be modified by the cell.

Evidence exists for both models. Treatment with doxorubicin decreases ecDNA copy number at the population level, with cells harboring higher copy numbers being more sensitive; this phenomenon is consistent with static selection^12^. By contrast, BET inhibitors can disrupt ecDNA tethering to chromatin, leading to mis-segregation of ecDNA during mitosis and global changes in copy number, consistent with active reconfiguration^13^. These seemingly contradictory results imply that ecDNA segregation may not always follow a purely stochastic, random process but can be contextually altered through specific molecular interactions. Nonetheless, a coherent framework explaining when and how these modes coexist is still lacking. Because selection of pre-existing variants and active reconfiguration are likely to operate together in a context-dependent manner, their interplay complicates the interpretation of ecDNA-mediated adaptation. Furthermore, conventional approaches for tracking ecDNA dynamics have limited throughput and cannot evaluate diverse stress contexts or clonal responses simultaneously^12,14–16^, making it difficult to achieve a comprehensive understanding of how ecDNA-mediated adaptation unfolds across heterogeneous populations.

To address these limitations, we established a high-throughput single-cell system that quantifies ecDNA copy-number distributions across thousands of cells and their derived lineages under defined perturbations. Using this approach, we asked whether cells can actively alter ecDNA segregation behavior and whether such alterations underlie adaptive evolution under drug-induced stress.

## Results

### Establishment of a high-throughput framework for the quantitative analysis of ecDNA copy-number heterogeneity

We developed a high-throughput single-cell DNA sequencing (scDNA-seq) framework based on the Tapestri platform, optimized for precise quantification of copy-number distributions across user-defined genomic regions. This integrative framework combines droplet-based scDNA-seq, cellular barcoding, and clonal tracking for quantitative, high-resolution monitoring of ecDNA dynamics across ∼11,000 individual cells and clonally related populations. We designed a fully customized primer panel that simultaneously amplifies target loci, control regions, and cellular barcode sequences, enabling both copy-number estimation and sample multiplexing. Multiplexing was achieved by combining two distinct cellular barcodes with a single nucleotide variant (SNV)-based indexing. To improve the accuracy of copy-number estimation, data from 40 evenly distributed control regions, obtained from both target cells and diploid RPE1 cells, were incorporated into a statistical normalization algorithm (**Fig. 1a** and **Extended Data Fig. 1a–c** and **2a–d**). Detailed methods are provided in the **Supplementary Note 1**.

**Fig. 1.**
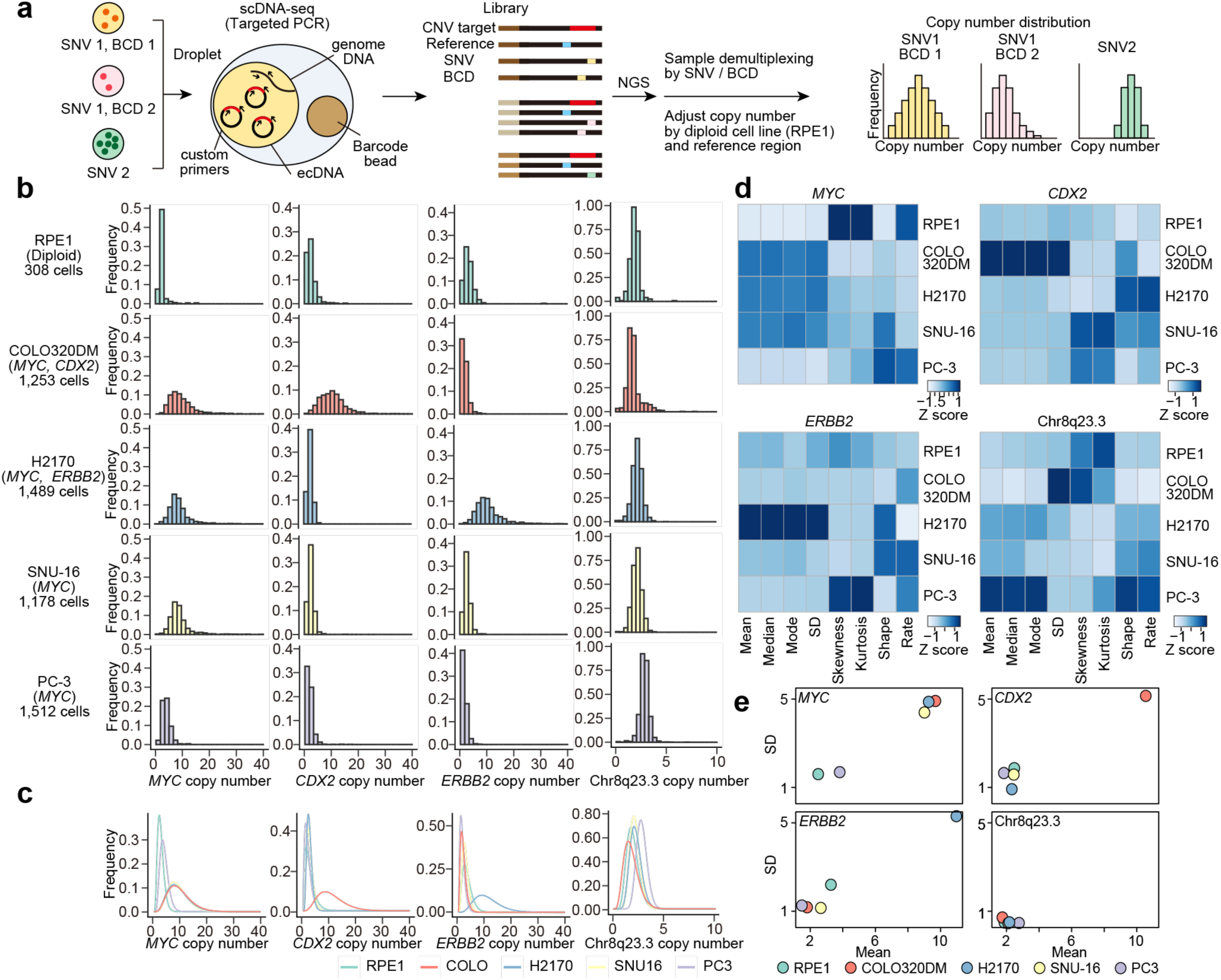
Establishment of high-throughput framework for quantifying extrachromosomal DNA (ecDNA) copy-number distribution. a, Schematic of the framework for the quantitative analysis of ecDNA copy-number distribution using single-cell DNA sequencing (scDNA-seq) based on the Tapestri platform. Droplet-based targeted PCR was used to quantify copy numbers of ecDNA-associated genomic regions and analyze their distributions. Single-nucleotide variations (SNVs) and cellular barcodes facilitated sample multiplexing and independent analysis within a single experiment, enabling a high-throughput workflow. **b,** Histograms of copy-number distributions of *MYC*, *CDX2*, *ERBB2*, and chr8q23.3 in ecDNA-positive cell lines (COLO320DM, H2170, SNU-16, and PC-3) and diploid cell line (RPE1). **c,** Gamma distribution fitting of copy-number distributions of each cell line shown in **b**. **d,** Heatmap of eight parameters for the cell lines shown in **b**. Each column was normalized by Z-score scaling. **e,** Scatter plot of the mean and standard deviation (SD) values showing simple and robust differences among cell lines.

This optimized system was used to analyzed the genomic regions known or predicted to reside on ecDNA across cancer cell lines harboring ecDNA, with the diploid cell line RPE1 serving as the control. Among the loci examined, *MYC* displayed broad and heterogeneous single-cell copy-number distributions in COLO320DM, H2170, and SNU-16 cells, whereas PC-3 cells showed only modest variability (**Fig. 1b**). Consistently, fluorescence in situ hybridization (FISH) confirmed the presence of ecDNA in the first three cell lines but not in PC-3, where all examined cells harbored only linear amplifications. Similarly, *CDX2* in COLO320DM and *ERBB2* in H2170 exhibited broad copy-number distributions, both of which have been shown to reside on ecDNA by FISH and previous reports (**Extended Data Fig. 2e**). These results indicate that even among amplified loci, those carried on ecDNA exhibit markedly greater copy-number variability, consistent with inherently uneven segregation during cell division.

Next, we evaluated how best to describe and parameterize their shapes across samples to establish a quantitative basis for comparing ecDNA copy-number distributions. The details of the statistical assessment are provided in **Supplementary Note 2** and **Extended Data Fig. 2d**. Based on these analyses, we defined three practical criteria for subsequent experiments as follows: (i) sampling at least 100 cells is sufficient to capture representative distributions; (ii) single-cell histograms can be fitted with gamma distribution facilitates visualization and parameter extraction; and (iii) the combination of mean and standard deviation (SD) provides a simple yet robust metric for comparative analyses. The copy-number distributions shown as histograms in **Fig. 1b** were fitted to gamma distributions, and eight candidate parameters were visualized as heatmaps together with a scatter plot of the mean and SD (**Fig. 1c–e**). This approach distinguishes cell lines not only by amplification status but also by the extent of heterogeneity within amplified regions, providing a quantitative framework for evaluating ecDNA copy-number variability at single-cell resolution.

### Clonal diversity of ecDNA copy-number distributions and transcriptional programs

To test whether ecDNA segregation follows a purely stochastic model, we examined how copy-number distributions evolve when starting from a single cell. Under stochastic and random segregation across all cells, independently derived clones are expected to exhibit similar binomially shaped distributions, even if their mean copy numbers differ due to initial copy numbers. Persistent interclonal differences, in contrast, would imply that segregation behavior is influenced by clone-specific mechanisms. To examine this, we performed single-cell cloning and then combined it with cellular barcode and scDNA-seq, allowing quantitative profiling of ecDNA copy-number distributions across multiple clones simultaneously (**Fig. 2a**).

**Fig. 2.**
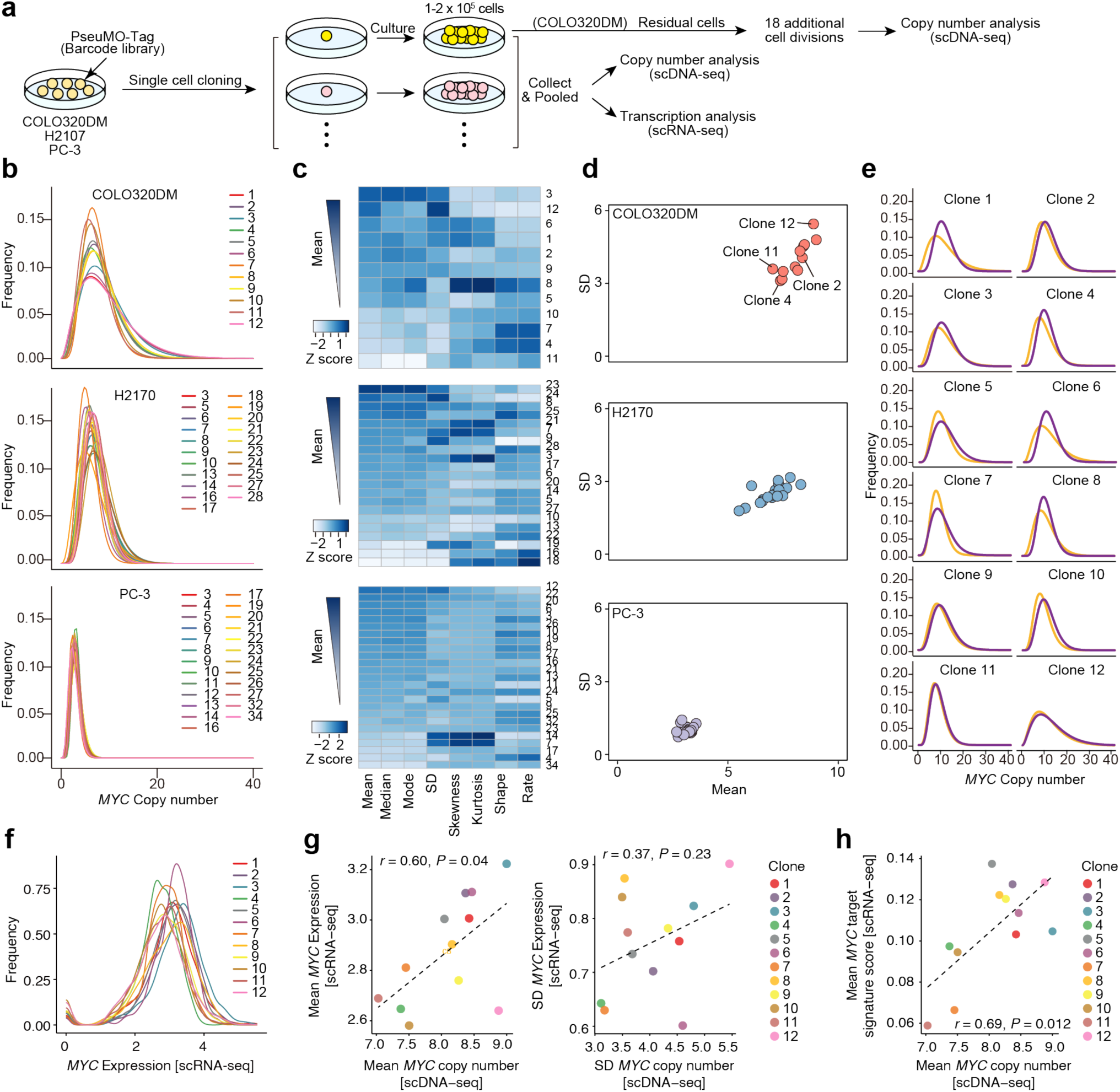
Clonal diversity of the uneven segregation patterns and transcriptional profiles. a, Schematic of single-cell cloning with cellular barcode. After PseuMO-Tag lentiviral cellular barcodes were transduced into each cell line, single cells were isolated into 96-well plates. Each clone was cultured until it reached approximately 1–2 × 10⁵ cells and cryopreserved. Once a sufficient number of clones was established, pooled clones were analyzed using scDNA-seq. For COLO320DM, the same clonal pool was additionally analyzed with scRNA-seq, and each clone was cultured individually. After 18 additional cell divisions, scDNA-seq was performed. **b,** Gamma distribution fitting of copy-number distributions of *MYC* for each clone derived from COLO320DM, H2170, and PC-3 cells. The dataset includes 5,861 cells from 12 clones in COLO320DM; 7,118 cells from 21 clones in H2170; and 14,796 cells from 25 clones in PC-3 cells. **c,** Heatmap of eight parameters of *MYC* for each clone shown in b, sorted in descending order of the mean value. Each column was normalized by Z-score scaling. **d,** Scatter plot of the mean and SD values for *MYC* copy number showing differences among clones. **e,** Gamma distribution fitting of *MYC* copy-number distributions in each COLO320DM clone. Orange curves represent the distributions at the same time point as in Fig. 2b**–d** and **Extended Data** Fig. 2. Purple curves represent the distributions after 18 additional cell divisions. **f,** Density plot of *MYC* expression levels for each clone. **g,** Correlation of the mean of *MYC* copy number and Mean *MYC* expression levels (left), and SD of *MYC* copy number and SD of *MYC* expression levels (right). **h,** Correlation of the mean of *MYC* copy number and mean of *MYC* target expression levels.

We applied this framework to three cancer cell lines—COLO320DM, H2170, and PC-3 — representing ecDNA-positive and ecDNA-negative contexts. In the ecDNA-positive lines, loci such as *MYC, CDX2*, and *ERBB2* exhibited broad single-cell copy-number distributions, consistent with the uneven segregation of ecDNA during mitosis (**Fig. 2b–d** and **Extended Data Fig. 3a–c**). By contrast, PC-3 cells displayed narrow distributions consistent with chromosome-integrated amplification (**Fig. 2b**). Notably, with each ecDNA-positive line, independently derived clones exhibited distinct copy-number distributions that deviated from the expectation of a uniform, random pattern. For example, in COLO320DM, some clones (e.g., clone 12) displayed broad distributions with large SD, whereas others (e.g., clone 2) maintained narrow distributions despite similarly high mean copy numbers. Clones 4 and 11 that had lower mean copy numbers also exhibited consistently narrow distributions (**Fig. 2c,d**). Similar interclonal differences were also detected for other ecDNA-associated genes — *CDX2* in COLO320DM and *ERBB2* in H2170 —whose copy numbers were correlated across clones (**Extended Data Fig. 3d**). We also confirmed high- and low-copy ecDNA states by FISH (**Extended Data Fig. 3e**). These observations indicate that, even under identical conditions, ecDNA copy-number distributions do not converge to a single steady-state form across clones, suggesting that not all clones undergo purely random segregation of ecDNA.

To determine whether these distributional differences are transient or stable properties of individual lineages, we further analyzed COLO320DM clones after approximately 18 additional cell divisions (**Fig. 2a**). Most clones retained the relative characteristics of their initial distributions, while some showed modest shifts in mean or variance. Clones that initially exhibited high copy numbers with large variability (clone 12), high copy numbers with low variability (clone 2), or low copy numbers with low variability (clones 4 and 11) showed little temporal change (**Fig. 2e** and **Extended Data Fig. 4a**), suggesting that their distinct segregation behaviors are self-maintaining over multiple generations. Furthermore, to identify transcriptional features associated with heterogeneity in ecDNA segregation among clones, we integrated scRNA-seq data from pooled COLO320DM clones collected at the same time point (**Fig. 2a** and **Extended Data Fig. 4b–d**). *MYC* expression levels varied among clones (**Fig. 2f**). Clone 2, which harbored a high copy number with low variability, displayed high and relatively uniform *MYC* expression. Clone 12, which carried an even higher copy number than clone 2, showed low *MYC* expression. Clones 4 and 11, which exhibited low copy numbers with limited variability through scDNA-seq, also showed lower *MYC* and *MYC*-target gene expression and reduced transcriptional variability in scRNA-seq (**Fig. 2g,h**). These results indicate that ecDNA copy-number state does not necessarily correlate with transcriptional output.

Together, these observations indicate that each clone maintains a distinct and stable ecDNA copy-number distribution and transcriptional output, implying that the mechanisms governing ecDNA maintenance differ among clones. Such diversity suggests that individual clones may rely on distinct ecDNA-associated regulatory programs, leading to heterogeneous adaptive responses under stress. To explore these possibilities, we selected four representative clones—2, 4, 11, and 12 —which exhibited stable yet divergent patterns in both segregation behavior and transcriptional profiles, for further investigation.

### Differential clonal responses to drug-induced ecDNA copy-number changes

We next asked whether these distinct ecDNA states translate into differential responses to environmental stress. To this end, we screened a panel of drugs and stress conditions to identify those in which ecDNA copy-number changes coincide with altered drug sensitivity. Among the conditions tested, the BET bromodomain inhibitor JQ1 and paclitaxel (PTX) induced significant reductions in *MYC* copy number, accompanied by enhanced survival specifically in the ecDNA-positive COLO320DM but not in the chromosomally amplified COLO320HSR (**Fig. 3a,b** and **Extended Data Fig. 5a-c**). These results suggest that under JQ1 and PTX treatment, changes in ecDNA copy number are associated with drug resistance.

**Fig. 3.**
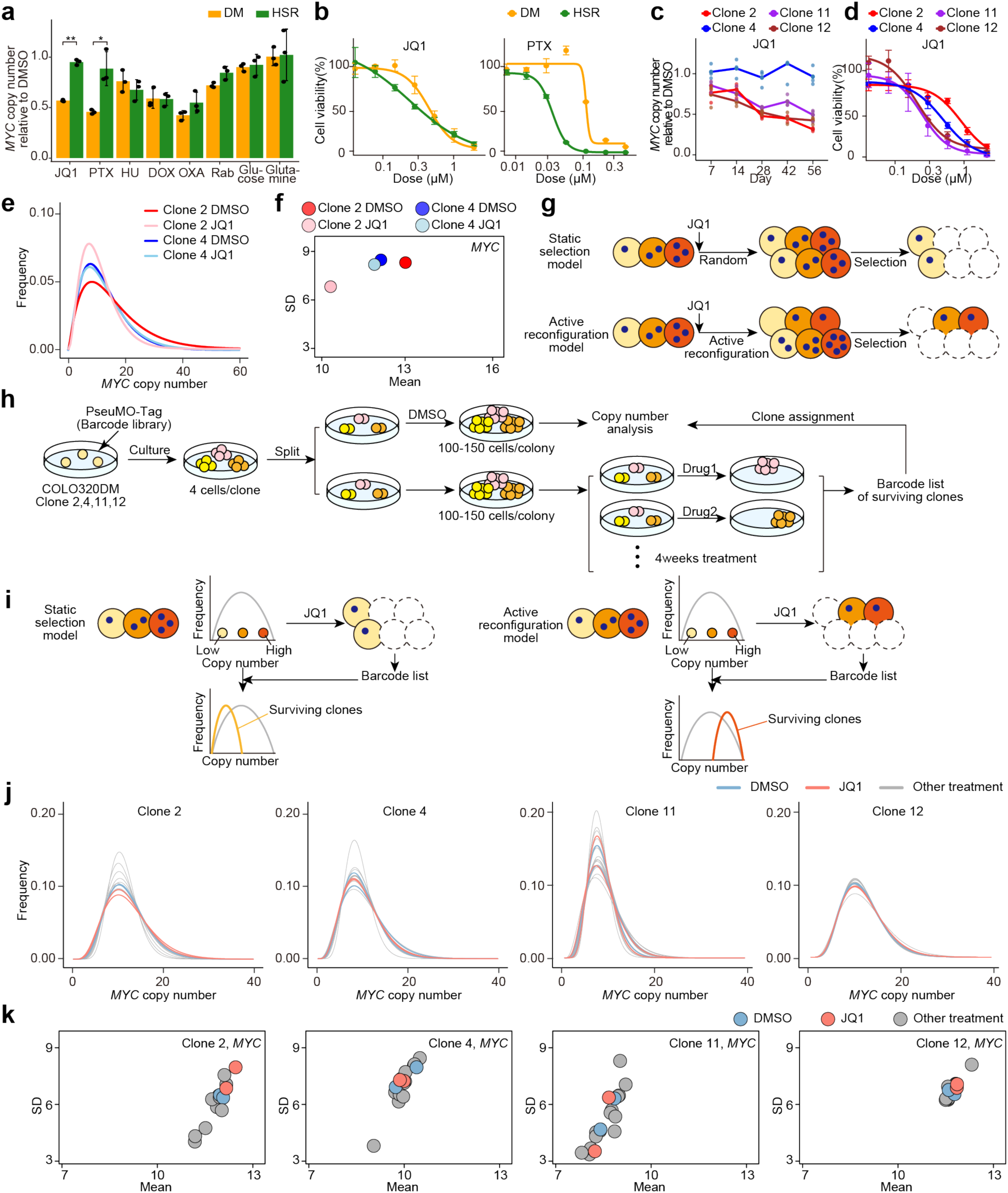
Clonal diversity in responses to drug-induced changes in ecDNA copy-number. a, *MYC* copy number of COLO320DM and COLO320HSR cells after 14-da treatment with various drugs or environmental stresses was measured by quantitative PCR and normalized to the DMSO-treated control of each cell line. Data are mean ± SD (n = 3). \*\**p* < 0.001 (JQ1), \**p* = 0.046 (PTX), *p* = 0.38 (HU, hydroxyurea), *p* = 0.95 (DOX, doxorubicin), *p* = 0.16 (OXA, oxaliplatin), *p* = 0.064 (Rab, rabusertib), *p* = 0.75 (Glucose; glucose depletion), and *p* = 0.92 (Glutamine; glutamine depletion). **b,** Cell viability assay of COLO320DM and COLO320HSR under the same conditions shown in a. Data are mean ± SD (n = 3). **c,** Time-dependent change in *MYC* copy number of COLO320DM clones 2, 4, 11, and 12 treated with 500 nM JQ1 over time, measured by qPCR and normalized to the DMSO-treated control of each clone. Each point represents an individual measurement, and the lines indicate the mean values (n = 3). **d,** Cell viability assay of JQ1-treated COLO320DM clones 2, 4, 11, and 12. Data are mean ± SD (n = 3). **e,** Gamma distribution fitting of copy-number distributions of *MYC* and chr8q23.3 in COLO320DM clones 2 and 4 treated with DMSO or JQ1 treatment. **f,** Scatter plot of the mean and SD values for *MYC* and chr8q23.3 copy number showing differences among conditions. **g,** Schematic of two hypothetical models for ecDNA behavior underlying survival under JQ1 treatment. **h,** Schematic of the experimental design for high-throughput clonal tracking using scDNA-seq and amplicon-seq under multiple stresses. **i,** Schematic of predicted experimental outcomes under the two hypotheses. **j,** Gamma distribution fitting of *MYC* copy-number distributions in each clone under drug-free conditions. The DMSO-treated group is shown in blue and the JQ1-surviving subclones in red. **k,** Scatter plot of the mean and SD values for *MYC* in each clone. The DMSO-treated group is shown in blue and the JQ1-subclones in red.

To determine whether the effects of JQ1 and PTX differ among ecDNA-positive clones, we monitored the time-dependent changes in *MYC* copy number and cell viability in the four representative COLO320DM clones (2, 4, 11, and 12). JQ1 and PTX treatment gradually reduced *MYC* copy number in clones 2, 11, and 12, but not in clone 4 (**Fig. 3c** and **Extended Data Fig. 5d**). Cell viability assays revealed that clone 2, which showed the greatest reduction in MYC copy number, was the most resistant to JQ1. Clone 12, which exhibited a high and variable ecDNA copy number, was more sensitive than clone 2. Clone 4, whose copy number remained largely unchanged, displayed an intermediate level of sensitivity (**Fig. 3d**) . These findings suggest that although JQ1 treatment reduces ecDNA copy number at population level, the sensitivity to JQ1 cannot be explained solely by differences in copy-number magnitude and heterogeneity. In contrast, under PTX treatment, clone 4 was the most resistant (**Extended Data Fig. 5e**). Taken together, these results indicate that the JQ1 resistance of clone 2 may be linked to alterations in ecDNA copy number.

To validate these observations at the single-cell level, we performed scDNA-seq on representative clones treated with JQ1. In clone 2, the copy-number distributions of *MYC* and *CDX2* — both ecDNA-associated loci, shifted leftward, showing decreases in both the mean and SD, whereas the *ERBB2* and chr8q23.3 regions remained unchanged. In clone 4, no appreciable changes were detected across any locus (**Fig. 3e,f** and **Extended Data Fig. 5f,g**). Consistent results were obtained in an independent experiment with shorter JQ1 exposure (7 days; **Extended Data Fig. 6a–c**), and FISH analysis after prolonged (56 days) treatment confirmed a reduction in ecDNA signals in clone 2 (**Extended Data Fig. 6d**). Together, these data confirm that JQ1 reduces ecDNA copy number at the single-cell level in a clone-specific manner.

A decrease in ecDNA copy number was observed, and two models could explain this phenomenon. The first is the static selection model, in which subclones with inherently low copy numbers preferentially survive JQ1 treatment. The second is the active reconfiguration model, in which subclones capable of generating low-copy cells, regardless of their original copy number, are the ones that survive (**Fig. 3g**). To determine which of these models is correct, we performed clonal tracking experiments. A lentiviral barcode library was introduced into each of the four representative COLO320DM clones (2, 4, 11, and 12), which were then evenly divide into two groups. One culture was analyzed by scDNA-seq to record baseline distributions, whereas the other culture was expanded, divided evenly, and exposed to various stresses for four weeks. Surviving clones under each condition were identified by barcode sequencing and linked to their corresponding scDNA-seq profiles, facilitating comparison of their pretreatment ecDNA states (**Fig. 3h**). If the static selection model is correct, JQ1- surviving clones should be those that already had low ecDNA copy numbers before treatment. In contrast, the active reconfiguration model predicts that surviving clones could originate from cells with either low or high pre-treatment copy numbers. Thus, if a JQ1- surviving clone originally had a low copy number, either model could explain its behavior; however, if the clone originally had a high copy number, this would indicate that active reconfiguration had occurred (**Fig. 3i**).

Data form qPCR and scDNA-seq analyses showed that clone 2 exhibited a marked reduction in copy number after JQ1 treatment. Interestingly, JQ1-surviving subclones derived from clone 2 exhibited higher and more heterogeneous MYC and CDX2 copy-number distributions at baseline (i.e., before JQ1 treatment) than subclones surviving under DMSO. These findings suggest that JQ1 resistance in clone 2 was not due to static selection of pre-existing low–copy-number cells, but rather involved active reconfiguration of ecDNA segregation. In contrast, PTX treatment also led to a reduction in copy number by qPCR, yet PTX-surviving subclones tended to originate from those with lower copy numbers at baseline, suggesting the occurrence of static selection in this context. Clone 4, which did not exhibit a copy-number change after JQ1 treatment, showed no association between ecDNA copy number and survival. Clone 11 and clone 12, both of which showed JQ1-induced copy-number reduction by qPCR, displayed patterns of surviving subclones distinct from clone 2. In clone 11, JQ1-surviving subclones tended to have lower baseline copy numbers, opposite to the trend in clone 2. In clone 12, the baseline copy-number distributions of surviving subclones were nearly identical across all stress conditions, indicating that ecDNA copy number was largely unrelated to survival in this clone (**Fig. 3j,k** and **Extended Data Fig. 7a,b**).

Taken together, these findings indicate that the mechanisms underlying drug resistance vary among clones and drug treatments. In particular, at least in clone 2, the reduction in ecDNA copy number upon JQ1 treatment is unlikely to result merely from static selection, but rather represents an active reconfiguration of ecDNA content.

### Clonal tracking reveals active reconfiguration of ecDNA in response to JQ1

To directly test whether JQ1-induced changes in ecDNA copy number reflected active reconfiguration rather than static selection, we conducted another clonal tracking experiment. A lentiviral barcode library was introduced into clone 2, which was then divided into two populations and treated with DMSO or JQ1 for 10 and 24 days, respectively, followed by scDNA-seq (**Fig. 4a**). If JQ1-surviving subclones also exhibit high ecDNA copy numbers under DMSO conditions, this suggest that active reconfiguration has occurred. The DMSO condition yielded 326 detectable subclones, whereas 119 subclones were recovered after JQ1 treatment (**Fig. 4b**). Consistent with the previous results, *MYC* and *CDX2* copy-number distributions shifted leftward after JQ1 exposure, whereas no changes were observed for the *ERBB2* and chr8q23.3 regions (**Extended Data Fig. 8a**).

**Fig. 4.**
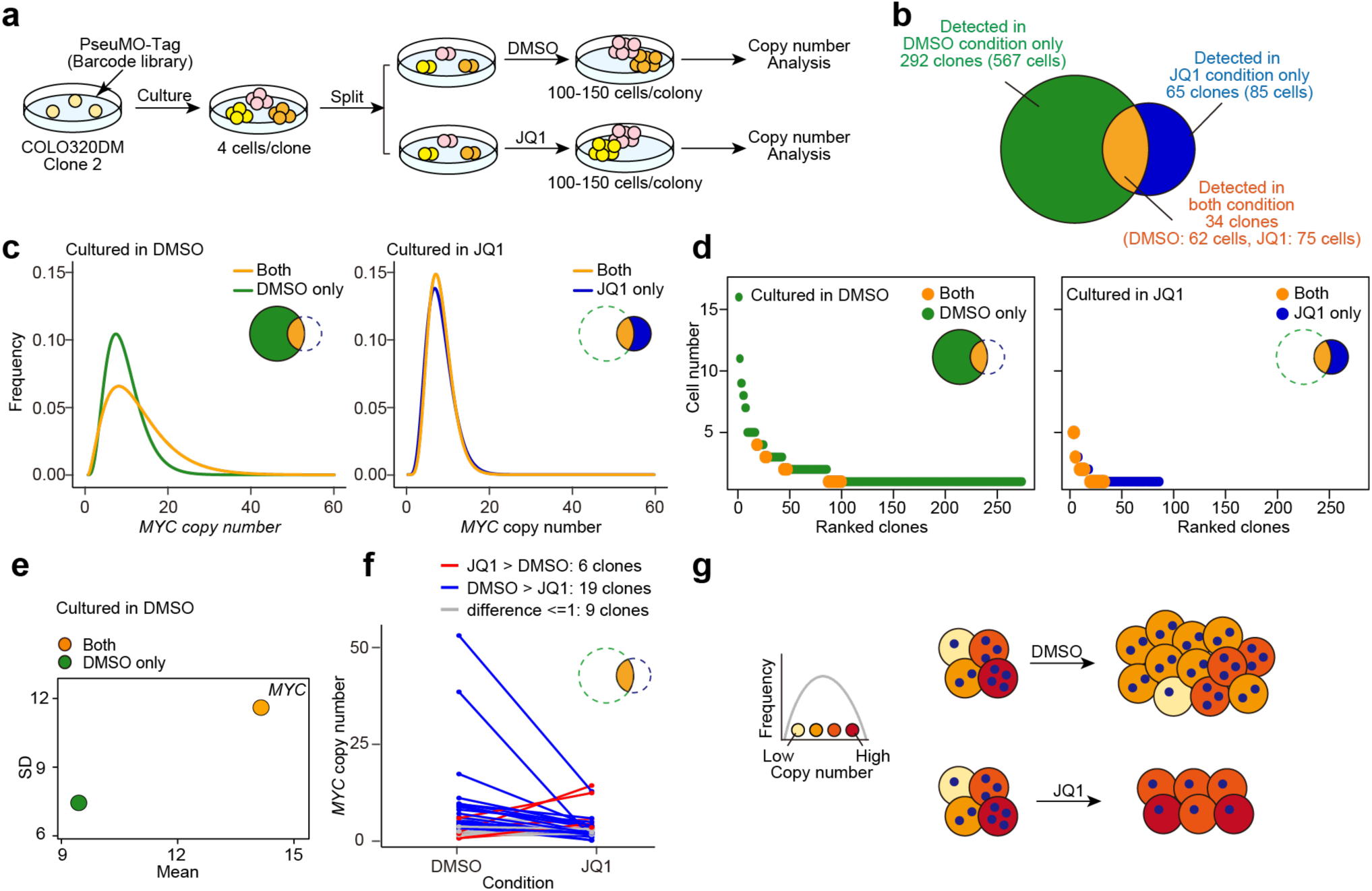
Active reconfiguration of ecDNA under JQ1 treatment revealed by clonal tracking. a, Schematic of the experimental design for clonal tracking using single-cell DNA sequencing (scDNA-seq) analysis in COLO320DM clone 2. One group was treated with DMSO and the other with JQ1. **b,** Venn diagram shows the number of cells and clones detected by scDNA-seq analysis in the DMSO-treated and JQ1-treated groups. **c,** The rank plot shows the number of cells detected per clone, sorted in the descending order of cell number. Clones detected in the DMSO-treated group are shown on the left, and those detected in the JQ1-treated group are on the right. Clones detected in both groups are highlighted in orange. **d,** Gamma distribution fitting of *MYC* copy-number distributions under DMSO (left) and JQ1 (right) conditions. Copy-number distributions of clones detected only in the DMSO-treated group are shown in green (left), those detected only in the JQ1-treated group are shown in blue (right), and detected in both groups are shown in orange. **e,** Scatter plot of the mean and SD values for *MYC* copy number in the DMSO-treated group. Clones detected only in the DMSO-treated group are shown in green, and those detected in both groups are shown in orange. **f,** Comparison of the mean *MYC* copy number between the DMSO and JQ1 culture conditions in clones detected in both the DMSO- and JQ1-treated groups. **g,** Schematic of cellular behavior under JQ1 treatment.

In the DMSO culture, approximately ten dominant subclones accounted for most of the population, indicating that selection had occurred even under drug-free conditions. Notably, the 34 subclones that survived JQ1 were not among these dominant subclones, implying that survival was not driven by pre-existing proliferative strength (**Fig. 4c**). Interestingly, under DMSO conditions, these JQ1-surviving subclones displayed higher and more heterogeneous *MYC* copy-number distributions, with greater mean and SD, compared with those that were lost under JQ1 (**Fig. 4d,e** and **Extended Data Fig. 8b,c**). Paired analyses of the same subclones without and with JQ1 exposure showed that *MYC* and *CDX2* copy numbers decreased following treatment (**Fig. 4f** and **Extended Data Fig. 8d**), excluding the possibility that survival arose from pre-existing low-copy states. Together, these results provide direct evidence that JQ1 response in clone 2 involves active reconfiguration of ecDNA content rather than static selection (**Fig. 4g**).

### Neuron-like transcriptional program distinguishes clone 2

To investigate the molecular basis underlying the distinct behavior of clone 2, we performed bulk RNA-seq on COLO320DM clones (2, 4, 11, and 12), their parental line, and COLO320HSR lacking ecDNA. Because transcriptional states associated with ecDNA are often linked to cell cycle–related genes, RNA was extracted under various culture and harvesting conditions that could potentially affect cell proliferation. Principal component analysis (PCA) revealed that the first three principal components (PC1, PC2, and PC3) captured the major sources of biological variability in the dataset (**Extended Data Fig. 9a**). PC1 reflected transcriptional differences between DM clones and HSR; PC2 captured variability between HSR and clone 4; and PC3 represented variability between HSR/clone 4 and the other clones (**Extended Data Fig. 9b–d**). These findings indicate that clones 2, 11, and 12 share similar transcriptional profiles, and that comparing clone 2 with clones 11 and 12 may help identify gene expression programs that are specifically enriched in clone 2. Differential expression analysis revealed 260 genes that were highly upregulated in clone 2 (**Extended Data Fig. 9e** and **Supplementary Table 4**). Notably, these activated genes were enriched for functions related to axon development and neuronal signal transduction (**Extended Data Fig. 9f**), suggesting that clone 2 possesses neuron-like transcriptional programs. The activation of neuronal-like pathways in clone 2 suggests that its microtubule dynamics and intracellular transport systems differ from those of typical cancer cells. During mitosis, such alterations could influence spindle assembly and chromosome– independent DNA partitioning, potentially affecting the segregation pattern of ecDNA. To test this possibility, we examined the sensitivity to JQ1 under nocodazole treatment which inhibits microtubule polymerization. When comparing JQ1 treatment alone with JQ1 combined with a low dose of nocodazole that does not affect cell proliferation, a marked synergistic effect of nocodazole was observed specifically in clone 2. In contrast, clone 12 showed no appreciable change in JQ1 sensitivity upon nocodazole addition (**Extended Data Fig. 9g,h**). Thus, the neuron-like transcriptional program of clone 2 may underlie a distinct mode of ecDNA redistribution under stress conditions such as BET inhibition.

## Discussion

In this study, we established a high-throughput and quantitative framework for measuring ecDNA copy-number distributions and their dynamics using scDNA-seq on the Tapestri platform. This approach enables integrated multi-omics analysis and clonal tracking with a cellular barcoding system. Using this approach, we revealed that certain cell populations actively alter the dynamics of ecDNA segregation in response to a specific stress, and that adaptive strategies differ across clones and stress conditions. In COLO320DM cells, bulk analysis showed that treatment with JQ1 reduces the copy number in certain clones. When we performed clonal tracking using two approachs within these clones —those already known to show reduced copy numbers in bulk—both experiments revealed that the decrease was not simply due to the pre-existence of low–copy-number cells that preferentially survived; rather, some clones survive by actively reconfiguring their segregation behavior in response to drug-induced stress. Notably, the clones capable of such reconfiguration exhibited neuron-like characteristics.

Methods for detecting ecDNA at the bulk level have rapidly advanced in recent years with the progress of genomic analysis technologies^17^. However, since ecDNA is known to exhibit copy-number heterogeneity, the development of single-cell–level analytical methods have been essential for investigating the biological significance of this heterogeneity. Although several image-based and plate-based methods have been developed, their throughput has remained limited. To overcome this limitation, we established a system based on scDNA-seq technology specifically optimized for analyzing ecDNA copy-number heterogeneity.

Although studies have reported that ecDNA copy number can change at the population level under drug stress, the underlying dynamics remain poorly understood. A recent study suggested that cells with high *MYC*-ecDNA copy numbers are more sensitive to doxorubicin^12^, whereas our results demonstrated that high-copy cells are not simply eliminated by JQ1 treatment; instead, they modulated their copy number to enhance survival. Moreover, doxorubicin sensitivity in our system was not directly associated with ecDNA copy number. Another recent study reported that JQ1 caused ecDNA loss through defective tethering in COLO320DM cells, and such a mechanism may contribute to the observed decrease in some clones^13^. Our finding that certain clones actively reconfigure their copy numbers is consistent with this model, yet our data further indicate that the mode of adaptation differs among clones. Importantly, our study also identified a distinct cellular phenotype associated with this reconfiguration: the cells exhibiting active reconfiguration displayed neuron-like characteristics, including unique microtubule organization and intracellular transport pathways. These features may contribute to the altered segregation behavior of ecDNA, distinguishing these cells from conventional cancer cells. This finding highlights that adaptation through ecDNA is intertwined with broader cellular state transitions, suggesting that genome architecture itself may serve as a flexible layer of non-genetic plasticity under stress.

A limitation of our study is that we did not experimentally dissect the molecular mechanisms of ecDNA segregation —for example, through gene knockout or perturbation experiments. Further studies will be required to understand the overall principles governing copy-number adaptation, our results reveal part of the underlying adaptive process.

Overall, our study establishes a high-throughput framework that enables quantitative analysis of ecDNA copy-number heterogeneity and dynamics at the single-cell and clonal levels. This approach provides a powerful tool to link the behavior of ecDNA to cellular adaptation. By demonstrating that ecDNA copy number is a dynamic and regulatable property rather than a static genomic feature, we uncover a new dimension of cellular adaptation to stress. This framework offers a foundation for future on how ecDNA-mediated genome plasticity contributes to tumor evolution, therapeutic resistance, and cellular adaptation.

## Methods

### Cell culture

COLO320DM and PC-3 cells were purchased from Japanese Collection of Research Bioresources (Osaka, Japan). H2170, SNU-16, and COLO320HSR cells were purchased from the American Type Culture Collection (ATCC, Manassas, VA). COLO320DM, COLO320HSR, and RPE1 cells were cultured in Dulbecco’s Modified Eagle’s Medium (DMEM) low glucose (Nacalai, Kyoto, Japan) supplemented with 10% FBS (Thermo Fisher Scientific, Waltham, MA). H2170, SNU-16, and SK-BR-3-Luc cells were cultured in Roswell Park Memorial Institute (RPMI, Nacalai) 1640 Medium supplemented with 10% FBS. PC-3 cells were cultured in Kaighn’s modification (F12K) of Ham’s F12 medium (Wako, Osaka, Japan) supplemented with 10% FBS. All cells were incubated at 37℃, 20% O2 and 5% CO2.

### Custom primer preparation for Single-cell DNA sequencing using the Tapestri Platform

Single-cell DNA analysis was conducted using the Tapestri Platform from Mission Bio (San Francisco, CA), which performs targeted PCR at the single-cell level. Custom primers were designed by Tapestri Designer (Mission Bio). We optimized the primers so that the single-nucleotide variant (SNV) region or cellular barcode locus required for cell-line or barcode identification was located within 75 bp from the reverse primer. We purchased oligonucleotides from Thermo Fisher Scientific, in which a bead-annealing constant sequence was appended to the 5′ end of the designed forward primer and the Nextera Read 2 sequence was appended to the 5′ end of the designed reverse primer. The constant sequence and Nextera Read 2 sequence appended were GTACTCGCAGTAGTC and GTCTCGTGGGCTCGGAGATGTGTATAAGAGACAG, respectively. For each run, the specific primers were mixed and diluted to 30 µM (forward primer mix) and 800 µM (reverse primer mix). To determine whether custom-designed primers could amplify target regions with efficiency comparable to that of commercially available primer panels, custom primers starting 1 bp upstream of the corresponding panel primers were mixed and evaluated in the same reaction. All custom primer sequences, excluding the common sequences added later, are listed in **Supplementary Tables 1.** and **2.** Supplementary Table 1. contains the primers used for the experiments, whereas Supplementary Table 2 includes the primers used for the validation of primer efficiency as described in Supplementary note 2.

### Single-cell DNA sequencing by Tapestri Platform

Single-cell DNA analysis was done using the Tapestri Platform with version 3 software and the reagents according to the version 3 user guide. Frozen cell stocks were thawed, washed once with phosphate-buffered saline (PBS), and passed through a 40-µm cell strainer. The cells were counted, mixed at various ratios of each analysis, and resuspended in Cell buffer at 3,000 cells/µL. They were loaded onto Tapestri Platform, followed by encapsulation, lysis, barcoding, and targeted PCR according the manufacturer’s instructions. Although the primers for the cancer-specific panels from Mission Bio were used as described in user’s guide, we primarily used the custom primers described above. Following targeted PCR, the emulsion was disrupted, the reaction was cleaned enzymatically, and the product was purified with AMPure XP beads (Beckman Coulter, Brea, CA). To add indexes, a second PCR of 10 cycles was performed, followed by AMPure XP beads (Beckman Coulter), and eluted in 15 µL. Library quality was assessed by Quantus (Promega, Madison, WI) and TapeStation 4200 (Agilent Technology, Santa Clara, CA) with D5000 Screen Tape. Next-generation sequencing (NGS) was performed in 2 x 75 bp paired-ends using NextSeq550 platform (Illumina, San Diego, CA, USA), which generate 50-60 million reads per library.

### Amplicon sequencing from a single-cell DNA sequencing library for cell barcode enrichment

NGS of the Tapestri library yielded few reads, including the cellular barcode locus, which rendered barcode assignment to individual cells difficult. This likely occurs because PCR efficiency at this locus was low and barcode-containing amplicons were under-represented. Therefore, a targeted enrichment PCR of barcode-containing amplicons was performed to increase representation and enable barcode assignment. Two cellular barcode systems, PseuMO-Tag and ID vector, were used. For PseuMO-Tag, the barcode-containing amplicon was shorter compared with the other amplicons, which resulted in failure to bind to AMPure XP beads (Beckman Coulter). Therefore, after targeted PCR and cleanup, the supernatant obtained from the first AMPure XP beads (Beckman Coulter) purification was used as a template for nested PCR. For the first PCR, 1 µL of the one-third–diluted supernatant was used as the template in a total reaction volume of 25 µL, using the NEBNext High-Fidelity 2× PCR Master Mix (New England Biolabs, Ipswich, MA). The PCR conditions were as follows: 30 sec denature step at 98℃, followed by seven cycles of 98℃ for 10 sec, 63℃ for 30 sec, and 72℃ for 30 sec, and finally 72℃ for 2 min. The primers used for the first PCR were as follows: Fw: 5’- CGTCGGCAGCGTCAGATG -3’, and Rv: 5’- GTCTCGTGGGCTCGGAGATGTGTATAAGAGACAGTCCGCTCGCTAGTTATTGCTCAACG -3’. To maintain barcode diversity, the reaction was initially run for seven cycles, then temporarily removed from the thermal cycler, and placed on ice. The appropriate number of additional PCR cycles was determined by qPCR. The qPCR reaction mixture (total 15 µL) contained 2 µL of the first PCR reaction as the template, 0.75 µL each of 10 µM forward and reverse primers, 0.24 µL of SYBR Green, 0.3 µL of ROX reference dye, 3.46 µL of nuclease-free water, and 7.5 µL of NEBNext High-Fidelity 2× PCR Master Mix. The amplification program was identical to that of the first PCR. The number of cycles corresponding to one-third to one-fourth of the plateau phase of the qPCR amplification curve was then added to the original reaction (kept on ice) to complete the first PCR. The PCR products were purified using 1.8x AMPure beads (Beckman Coulter), and eluted in 17 µL. For the second PCR, 1 µL of the first PCR product template in total amount of 25 µL reaction volume was used with NEBNext High-Fidelity 2x PCR Master Mix (New England Biolabs). The PCR conditions were as follows: 30 sec denature step at 98℃, followed by 5 cycles of 98℃ for 10 sec, 61℃ for 30 sec, and 72℃ for 30 sec, and finally 72℃ for 2 min. The primers used in the second PCR were as follows: Fw: 5’- AATGATACGGCGACCACCGAGATCTACACXXXXXXXXTCGTCGGCAGCGTCAGATG -3’, and Rv: 5’- CAAGCAGAAGACGGCATACGAGATXXXXXXXXGTCTCGTGGGCTCGGAGA -3’. X indicates the indices. The PCR products were purified using 0.8x→1.3x SPRIselect (Beckman Coulter), and eluted in 17 µL. For the ID vector, PCR was carried out under the same conditions. The primers used for the first PCR were as follows: Fw: 5’- CGTCGGCAGCGTCAGATG -3’, and Rv: 5’- GTCTCGTGGGCTCGGAGATGTGTATAAGAGACAGCGAGGCAGGAAACAGTGACTAG -3’. After the initial seven cycles, the reaction was placed on ice, and qPCR was done to determine the appropriate number of additional cycles, based on the same procedure used for the PseuMO-Tag amplicon. After purification with 1.8x AMPure beads (Beckman Coulter), the second PCR was performed using the same primers and PCR conditions were similar to those used for the PseuMO-Tag amplicon. The PCR products were purified using 0.7x SPRIselect (Beckman Coulter), and eluted in 17 µL. Library quality was assessed by Quantus (Promega, Madison, WI) and TapeStation 4200 (Agilent Technology) with D5000 Screen Tape. The expected amplicon size was 256 bp for the PseuMO-Tag and 374–382 bp for the ID vector. NGS was performed in 2 x 75 bp paired-ends using NextSeq550 platform (Illumina), which generates 5 million reads per library.

### Single-cell DNA sequencing data processing pipeline and normalization

A custom computational pipeline was established to construct copy-number count matrices and to annotate cell identities according to SNV profiles (see *Code availability*). For each single-cell DNA sequencing (scDNA-seq) dataset, paired-end FASTQ files were processed by the extracting sequence reads and concatenating Read1 and Read2 using a tab delimiter, followed by sorting before inputting into the pipeline. Read1 consisted of the Tapestri barcode (TapeBC) and forward primer sequence, whereas Read2 contained the reverse primer and downstream genomic sequence. Based on the primer, Read2 sequences corresponded to copy number variant quantification, SNV detection, reference region, or cellular barcodes (PseuMO-Tag and ID vector). Within the pipeline, each read was scanned to extract the TapeBC. The reads sharing identical barcodes were aggregated, and per-target read counts were calculated to generate the copy-number matrix. The reads containing SNV information were then used to separately count wild-type and mutant alleles, thus producing an SNV count matrix from which allele frequencies (AF) were determined. Based on the AF thresholds, the cells were classified as wild-type (AF > 0.8), heterozygous (0.2 ≤ AF ≤ 0.8), or mutant/LOH (AF > 0.8 for mutant reads). These SNV patterns were compared with known mutation profiles for each cell line to annotate the corresponding cell identity, which was linked to the copy-number count matrix. Raw read counts were normalized based on reference regions with low variability (top 20 regions with the smallest coefficient of variation in RPE1 cells). For each cell, the mean count across the reference regions was used for depth normalization. Correction factors were then calculated for each target region by trimming extreme values and aligning the modal copy number of the RPE1 cells to two. Normalized and adjusted copy numbers were then calculated for all cells and exported for downstream analysis.

### Statistical analysis and visualization of copy number

Density-scaled histograms were generated to visualize the distribution of copy numbers for each cell line. The cell lines harboring ecDNA frequently showed right-skewed distributions with extended tails. To characterize these distributions, the copy-number distributions were also visualized using the fitted gamma curves (see ***Supplementary Note 2***). As quantitative indicators of the distribution shape, the mean, median, and mode (representing copy-number level), SD (representing dispersion), skewness (asymmetry), kurtosis (peakedness), and gamma distribution parameters, shape and rate were calculated. These parameters were visualized using heatmaps and scatter plots.

### Single-cell DNA sequencing –PseuMO-Tag assignment

The clone-specific barcode sequences obtained from cDNA amplicon sequencing (amplicon-seq) were initially compiled as a whitelist (*see Preparation of amplicon-seq library from cDNA Section*). Based on the amplicon-seq FASTQ files enriched from the scDNA-seq library, the TapeBC and PseuMO-Tag sequence regions were extracted using cutadapt (Hamming distance ≤ 1). For each TapeBC, the read counts were aggregated across the clones, and the proportion of reads assigned to each clone was determined. The cells in which > 80% of the reads mapped to a single clone were designated as clone-assigned. The clone labels were merged with the normalized copy-number matrix for downstream analyses. Statistical analyses and data visualizations were similarly conducted on a per-clone basis, following the same procedures described above.

### Single-cell DNA sequencing – ID assignment

ID-specific barcode sequences obtained from Sanger sequencing were first compiled as a whitelist (see *Sanger sequencing section*). The subsequent procedures for barcode extraction, read-count aggregation, clone assignment, and integration with the copy-number matrix were conducted in the same manner as described above.

### Single-cell RNA sequencing analysis

Single cell RNA sequencing data analysis and clone assignment were performed as previously reported^18^.

### DNA FISH probe preparation

Bacterial artificial chromosome (BAC) clones in *E. coli* were purchased from Advanced GenoTechs (Ibaraki, Japan) and cultured in 200 mL of lysogeny broth (LB) medium containing 34 µg/mL chloramphenicol. The following day, BAC clones were extracted using QIAGEN Plasmid Maxi Kit (QIAGEN, Gelderland, Netherlands). Probes were enzymatically labeled at 15℃ for 12h using Nick translation kit (Abbott, IL, USA) with either Fluorescein-12-dUTP (Roche, Switzerland) or ChromaTide Texas Red-12-dUTP (Thermo Fisher Scientific). The following day, labeled oligos were purified using MicroSpin G50 Columns (Roche, Switzerland) according to the manufacturer’s instructions. Briefly, columns were centrifuged at 740 x g for 1 min to remove storage buffer, samples were applied to the columns, and probes were recovered by centrifugation. The following BAC clones were used: RP11-1136L8(*MYC*) and RP11-94L15(*ERBB2*).

### Metaphase chromosome spread

For, COLO320DM and COLO320HSR, cells in the metaphase were collected following the treatment with KaryoMAX (Thermo Fisher Scientific) at 100 ng/mL for 6 h. For other cell lines, cells in the metaphase were collected following the treatment with Colcemid (AdipoGen, Switzerland) at 100 ng/mL for 6 h. All cell lines were washed once with PBS, and single-cell suspensions were incubated under hypotonic condition (3:7 PBS: deionized distilled water) for 5min at room temperature. The samples were fixed in Carnoy’s fixative (3:1 methanol: acetic acid), and the cell pellet were collected by concentration. Samples were fixed additional three times with Carnoy’s fixative. Finally, they were resuspended in the fixative and dropped onto MAS-coated slide glass (Matsunami, MAS-01, Osaka, Japan).

### Metaphase DNA FISH

Fixed cells on slide glasses were air-dried overnight. The hybridization mix was prepared by combining 50 µL of deionized formamide (Sigma-Aldrich, St. Louis, MO), 10 µL of 20× SSC (Sigma-Aldrich), 20 µL of 60% dextran sulfate (Wako), 10 µL of water, and 1 µL of Tween 20 (Wako), and thoroughly mixed. Next, the probe mix was prepared. For each slide, 10 µL of red-labeled probe, 10 µL of green-labeled probe, 2.5 µg of Human Cot-1 DNA (Thermo Fisher Scientific), 10 µg of sonicated salmon sperm DNA (BioDynamics Laboratory), 2.6 µL of 3 M Na₂CO₃ (Nacalai), and 65 µL of ice-cold 100% ethanol were mixed and centrifuged at 15,000 rpm for 5 min at 4 ℃. The supernatant was discarded, and the pellet was retained. Next, 10 µL of the hybridization mix was added to the pellet and mixed well, and the entire mixture was dropped onto a glass slide. A coverslip (Matsunami) was placed on top, sealed with paper bond (Kokuyo, Osaka, Japan), denatured at 80 ℃ for 1 min, and immediately incubated at 37℃ in a humidified light-protected chamber. The following day, the coverslip was removed, and the slides were washed twice with PBS. The samples were stained with 4’,6-diamidino-2-phenylindole (DAPI) (1:1000 in PBS) for 5 min and mounted with 10 µL of ProLong Gold with DAPI (Thermo Fisher Scientific). Images were captured using BZ-X700 microscope (Keyence, Osaka, Japan) with a 100× objective lens.

### Preparation of PseuMO-Tag lentiviruses

PseuMO-Tag lentiviruses were prepared as previously reported^18^. Briefly, the construct was derived from the pLenti-CMV-GFP-Puro vector (Addgene #17448), with a strong CMV promoter immediately upstream of the barcode sequence. Barcode sequence was inserted by Golden Gate assembly. Lentiviruses were produced using FuGENE HD Transfection Reagent (Promega).

### ID vector construction

The construct was modified based on the pLenti-CMV-GFP-Puro vector (Addgene #17448), with a strong CMV promoter immediately upstream of the barcode sequence. The original vector was digested with *EcoRI* and *KpnI* to remove the PGK promoter and puromycin resistance gene, and replaced with the EF-1α core promoter and an EGFP-P2A-puromycin resistance cassette. The vector was then digested with *ClaI* and *SalI* to remove the original CMV promoter and EGFP, and a new CMV promoter, followed by a stuffer sequence for the cellular barcode, was inserted. The resulting plasmid served as the backbone for the cellular barcode vector. Next, 10 µg of the plasmid was digested with *BsmBI-v2*, purified using the QIAquick Gel Extraction Kit (QIAGEN), and eluted in 20 µL of elution buffer. For insertion, 2 µL of 100 µM barcode oligonucleotide and 1 µL of 100 µM reverse primer (5′-TGCAGCATGCGTCTCACAACG-3′) were mixed with 25 µL of NEBNext High-Fidelity 2× PCR Master Mix (New England Biolabs) and 22 µL of water to produce double-stranded barcode DNA (Thermo Fisher Scientific). PCR was carried out using the following conditions: 98 °C for 2 min; 10 cycles of 65 °C for 30 sec and 72 °C for 10 sec; and a final extension at 72 °C for 2 min. The PCR product was purified using a QIAquick PCR purification kit (QIAGEN) and eluted in 20 µL.

The backbone plasmid (1.5 µg) and barcode insert (33 ng) were ligated by Golden Gate assembly. The reaction mixture consisted of 2 µL of r3.1 buffer (New England Biolabs), 1 µL of *BsmBI-v2* (New England Biolabs), 1 µL of T4 DNA ligase (New England Biolabs), 2 µL of 10× T4 DNA ligase buffer (New England Biolabs), and nuclease-free water in a total volume of 20 µL. The reaction was incubated in a thermal cycler under the following conditions: 100 cycles of 42 °C for 2 min and 16 °C for 5 min, followed by incubation at 50 °C for 10 min and 80 °C for 20 min. The reaction product was purified using DNA Clean & Concentrator Kit (Zymo Research, Irvine, CA) and eluted in 10 µL. The purified plasmid was transformed into NEB Stable Competent *E. coli* (New England Biolabs), which were cultured in 500 mL LB medium containing 100 µg/mL ampicillin at 30 °C overnight with shalking. Plasmid DNA was then extracted using QIAGEN Plasmid Plus Maxi Kit (QIAGEN). To isolate individual barcode plasmids from the mixed-barcode plasmid pool, the mixture was transformed into NEB Stable Competent *E. coli* (New England Biolabs). The following day, 20 colonies were randomly selected and cultured in 3 mL of LB medium at 37 °C with shaking overnight. Plasmid DNA was purified using the Monarch Plasmid Miniprep Kit (New England Biolabs).

### Sanger sequencing

The barcode sequence for each plasmid was validated by Sanger sequencing. The reaction mixture (10 µL total volume) consisted of 150–300 ng of plasmid DNA, 1 µL of 3.2 µM primer, 2 µL of BigDye Terminator v3.1 (Thermo Fisher Scientific), 1 µL of 5× sequencing buffer, and nuclease-free water. The reactions were incubated as follows: 96 °C for 1 min, followed by 25 cycles of 96 °C for 10 sec, 50 °C for 3 sec, and finally 60 °C for 75 sec. The sequencing primer used was 5′- TAAGCAGAGCTATGGTGAGC-3′. The products were purified using a Gel Filtration Cartridge (Edge BioSystems, San Jose, CA) and transferred to a MicroAmp Optical 96-Well Reaction Plate (Thermo Fisher Scientific), denatured at 94 °C for 2 min, and subjected to capillary electrophoresis on a 3500xL Genetic Analyzer (Thermo Fisher Scientific).

### ID vector transfection

For each well of a 96-well plate, 5 µL of Opti-MEM (Thermo Fischer Scientific) and 0.32 µL of FuGENE HD Transfection Reagent (Promega) were mixed and incubated at room temperature for 5 min. The mixture was then added to 80 ng of ID vector that was pre-aliquoted into another tube, followed by incubation at room temperature for 15 min. Next, 45 µL of the culture medium was added, and the mixture was gently pipetted twice. Finally, 50 µL of medium was removed from each well, and the prepared transfection mixture was added.

### Reagents and special medium

JQ1, paclitaxel, oxaliplatin, doxorubicin, and hydroxyurea were purchased from Selleck Biotech (Kanagawa, Japan). Glutamine depleted DMEM and glucose depleted DMEM were purchased from Nacalai. Medium containing drugs was replaced every 3-4 days.

### RNA extraction and reverse transcription

Total RNA was extracted using RNeasy Mini/Micro Kit (QIAGEN) according to the manufacturer’s instructions. Reverse transcription was performed with 10 ng of RNAs using PrimerScript RT Master Mix (TAKARA, Shiga, Japan).

### Preparation of amplicon-seq library from cDNA

To prepare a list of PseuMO-Tag sequences for each clone, amplicon-seq for the barcode locus was performed. The library was prepared using nested PCR as previously reported^18^. For the first PCR, 1 µL of the cDNA template in a total volume of 25 µL was used with NEBNext High-Fidelity 2x PCR Master Mix (New England Biolabs). The PCR conditions were as follows: 30 sec denature step at 98℃, followed by 35 cycles of 98℃ for 10 sec, 67℃ for 30 sec, and 72℃ for 30 sec, and finally 72℃ for 2 min. The primers used for the first PCR were as follows: Fw: 5’- CACCATCGTGGAACAGTACG -3’, and Rv: 5’- GTCCGCTCGCTAGTTATTGC -3’. The PCR products were purified using 0.6x SPRIselect beads (Beckman Coulter), and eluted in 12 µL. For the second PCR, 1 µL of the first PCR product template in a total volume of 50 µL was used with NEBNext High-Fidelity 2x PCR Master Mix (New England Biolabs). The PCR conditions were as follows: 30 sec denature step at 98℃, followed by 8 cycles of 98℃ for 10 sec, 67℃ for 30 sec, and 72℃ for 30 sec, and finally 72℃ for 2 min. The primers used for the second PCR were as follows: Fw: 5’- AATGATACGGCGACCACCGAGATCTACACXXXXXXXXACACTCTTTCCCTACACGACGCTCTTC CGATCTGCAGGAAACAGCTATGACTATGC -3’, and Rv: 5’- CAAGCAGAAGACGGCATACGAGATXXXXXXXXGTGACTGGAGTTCAGACGTGTGCTCTTCCGA TCTGTCCGCTCGCTAGTTATTGC -3’. X indicates the indices. The PCR products were purified using 0.8x→1.2x SPRIselect (Beckman Coulter), and eluted in 12 µL. Library quality was assessed by Quantus (Promega) and TapeStation 4200 (Agilent Technology) with D1000 Screen Tape. NGS was performed in 2 x 38 bp paired-ends using NextSeq550 platform (Illumina), which generates 0.5-1 million reads per library.

### DNA extraction

Quantitative PCR (qPCR) was used to evaluate changes in ecDNA copy number following drug treatment. DNA was extracted from cells following lysis. The cell pellets were collected and resuspended in 36 µL of 50 mM NaOH, followed by incubation at 95 °C for 10 min. After cooling to room temperature, 4 µL of 1 M Tris-HCl (pH 8.0) was added to neutralize the solution. The mixture was then diluted with 120 µL of nuclease-free water. NaOH solution was freshly prepared by appropriate by diluting from a 1 M stock. For other experimental purposes, DNA was extracted using QIAmp UCP DNA Micro Kit (QIAGEN) or DNeasy Blood & Tissue Kit (QIAGEN) according to the manufacturer’s instructions.

### Quantitative PCR

Quantitative PCR was done using TB Green® Premix Ex Taq™ II (Tli RNaseH Plus) (TAKARA) and StepOnePlus (Thermo Fischer Scientific), according to the manufacturer’s instructions. To determine the internal control, as shown in Extended Data Fig. 5A, 0.6 ng of DNA was used as a template. To evaluate copy number changes following drug treatment, 2 µL of the lysate-extracted DNA described above was used as the template per well. The internal control was chr5q23.1, which showed no difference in amplification compared with RPE1 (**Extended Data Fig. 5a**). Amplification levels were normalized relative to chr5q23.1 levels, using the ΔΔCT method. The primers were as follows: Chr4-Fw 5’- CATGCTCTATCTACACTTGTAAGCCA -3’, Chr4-Rv 5’- GCCCTGCAGCCTAGGAATG -3’, Chr5q23.1-Fw 5’- TTTGTTCTACTACAAAGACTCATGCAC -3’, Chr5q23.1-Rv 5’- CTGGGTTGGTTCCAGGTCTT -3’, Chr6-Fw 5’- AGAAGGGATTACACTATCTGTCACTC -3’, Chr6- Rv 5’- AACCACCATACCCAGCCATG -3’, Chr17-Fw 5’- GCCAGACCTGTGAATATTTTATGTTAC - 3’, Chr17-Rv 5’- TTCTTCTCCCTCCCTCGTCC -3, ’Chr22-Fw 5’- ACCCTCCTATTAAAATGCCTCTGA -3’, Chr22-Rv 5’- ACGGCGCATGAACAGAAAAC -3’, MYC-Fw 5’- GGACGACGAGACCTTCATCA -3’, MYC-Rv 5’- GAGGCCAGCTTCTCTGAGAC -3’, and ERBB2-Fw 5’- GCTGGTACTTTGAGCCTTCACA -3’, ERBB2-Rv 5’- GAAGGCGGGAGACATATGGG -3’.

### Preparation of amplicon-seq library from DNA

To preserve barcode diversity, 50 ng of DNA was used as the template. To maintain barcode diversity, the first PCR was performed for seven cycles under the same temperature conditions as described above. Subsequent purification and the second PCR were performed as described above. NGS was performed in 2 x 38 bp paired-ends using NextSeq550 platform (Illumina), which generates 2-5 million reads per library.

### Cell viability assay

COLO320DM and COLO320HSR were seeded into 96-well plates at 2,000 cells/well with three replicates, and the drugs were added the following day. After 7 days, 10 µL of CCK-8 reagent from Cell Counting Kit-8 (Dojindo, Kumamoto, Japan) was added to each well, and then the absorbance at 450 nm was measured using Bio Tek Gen5 (Agilent Technology). The absorbance was normalized to that of DMSO-treated wells.

### Cell proliferation assay

Cell proliferation was measured using the IncuCyte S3 Live-Cell Analysis System (Sartorius, Germany). Similar to the cell viability assay, cells were seeded into 96-well plates at 2,000 cells per well in triplicate, and drugs were added the following day. Confluency values were measured at 0 and 144 hours, and growth rates were calculated and compared across conditions.

### Statistical analyses

Two-sample comparisons were performed using a two-sided Welch’s t-test. For all analyses, statistical significance was set at *p* < 0.05.

### Bulk RNA sequencing library preparation

Total RNA extracted with RNeasy Plus Mini Kit (QIAGEN). Each library was constructed from 1 µg of RNA using the SMARTer Stranded Total RNA Sample Prep Kit - HI Mammalian (TAKARA), according to the manufacturer’s instructions. Library quality was assessed by Quantus (Promega) and TapeStation 4200 (Agilent Technology) with D5000 Screen Tape. Next generation sequencing was performed in 2 x 38 bp paired-ends using NextSeq550 platform (Illumina), which generates 20 million reads per library.

### Bulk RNA sequencing data analysis

RNA sequencing data were analyzed as previously reported^19^. Briefly, the row reads were trimmed by Skewer (v0.2.2), followed by mapping to GRCh38 genome by STAR (v.2.7.8a), and then counted by featureCounts (v.2.0.10). Differential gene expression analysis was performed with edgeR’s glmQLFTest (v3.32.1). Gene Ontology enrichment analysis was performed by ClusterProfiler’s enrichGO() function.

### Clonal tracking by scDNA-seq and amplicon-seq (Figure 3)

COLO320DM Clone 2, Clone 4, Clone 11, and Clone 12 cells were seeded into 96-well plates at 10,000 cells per well. The following day, cells were transduced with PseuMO-Tag lentiviruses (MOI = 3) supplemented with 10 µg/mL polybrene. One day after transduction, cells were trypsinized, dissociated, and resuspended in 20 mL of DMEM. The suspension was distributed into a new 96-well plate at 150 µL per well (Split 1). When the colonies reached 4–8 cells per clone (4 days after plating for all clones), the number of clones containing ≥ 4 cells per clone was counted. The cells from each well were then trypsinized again to obtain single-cell suspensions. Wells were pooled so that each mixture contained approximately 100–120 clones, and the mixed suspension was equally divided into two wells of a new 96-well plate (Split 2). The following day, one well of each pair was treated with 1:1000 DMSO, and the other was maintained without any treatment. For the DMSO-treated group, the cells were cultured until the colonies reached 100–150 cells per colony (Clone 2, 9 days; Clone 4, 9 days; Clone 11, 14 days; Clone 12, 8 days). Next, the ID vector was transfected as described above, and the cells were collected 18–20 h later. To obtain a total of approximately 600 clones, cells from six corresponding well pairs were pooled, generating six mixed samples for storage. The clones remained spatially separated throughout the culture period, without any physical contact or competition. After Clone 11 collection, all four clones were pooled and subjected to scDNA-seq. For the untreated paired wells, six corresponding wells were harvested on the same day as the DMSO group, pooled, and split into 18 wells of a 96-well plate (Split 3). The following day, the cultures were treated with 1:1000 DMSO, 500 nM JQ1, 100 nM paclitaxel, 50 µM hydroxyurea, 300 nM doxorubicin, 10 µM oxaliplatin, and 500 nM rabusertib, glucose depletion, or glutamine depletion. The medium was replaced every 3–4 days. After four weeks, the cells were collected for DNA extraction and amplicon-seq library preparation (see *DNA extraction Section and Preparation of amplicon-seq from DNA Section*). Based on the amplicon-seq results, the clones with barcodes detected by two or more sequencing reads were defined as surviving clones under each stress condition. These surviving clones were associated with the scDNA-seq data. For each clone, the mean copy number across all targeted regions was determined and used for comparative statistical analyses of copy-number distributions.

### Clonal tracking by scDNA-seq (Figure 4)

COLO320DM Clone 2 cells were seeded into 96-well plates at 10,000 cells per well. The following day, cells were transduced with PseuMO-Tag lentiviruses (MOI = 3) supplemented with 10 µg/mL polybrene. The day after transduction, the cells were trypsinized, dissociate, and resuspended in 10 mL of DMEM. The suspension was distributed into a new 96-well plate as follows: (i) 100 µL per well for 48 wells, and (ii) 60 µL per well for additional 48 wells (Split 1). Each well represented an independent clone that could be tracked using a unique combination of barcodes. When the colonies reached 4–8 cells per clone (4 days after plating), the number of clones containing ≥ 4 cells was counted. Cells were again trypsinized to obtain single-cell suspensions. To prepare mixed populations containing approximately 50–60 clones, one well from group (i) and one well from group (ii) were combined, and equally divided into two new wells of a 96-well plate. This procedure was repeated for all pairs, resulting in 48 paired wells, each containing 50–60 mixed clones (Split 2). The following day, one well for each pair was treated with 500 nM JQ1, whereas the other was treated with DMSO as a control. Cells were cultured until colonies reached 100–150 cells per colony (10 days for the DMSO group and 24 days for the JQ1 group). At that point, the ID vector was transfected to label each condition, and cells were collected 18–20 h after transfection. As the goal was to analyze approximately 300 clones, cells from six corresponding well pairs were pooled, generating six mixed wells for storage. The transfection procedure followed the protocol described below (see *ID vector transfection section)*. Throughout the culture period, individual clones remained spatially separated, with no physical contact or competition between the clones. After collection, samples from the JQ1 and DMSO groups were pooled for scDNA-seq. During the analysis, clones detected in the DMSO and JQ1 groups were extracted separately. Of these, we focused on two subsets: clones detected only in the DMSO group, but not in the JQ1 group, and those detected in both the DMSO and JQ1 groups. The differences in their copy-number distributions were then analyzed and compared.

## Data availability

Data associated with this work is available at BioStudies. accession E-MTAB-16119.

## Code availability

Processed single-cell objects and custom PseuMO-Tag Decoder to reproduce analyses and figures is available at https://github.com/Chikako-Shibata/ecDNA_single_cell_DNA.

## Acknowledgements

We gratefully acknowledge the technical assistance of Mana Nagamoto, Kaoru Masuda, Hiromi Ayame, Noriko Kaneniwa, Hiromi Otsuka, and Nami Nakasuji in experiments and data analysis. We also thank all members of our laboratory for valuable discussions and continuous encouragement. English language editing was provided by Enago (www.enago.jp).

## Author contributions

C.S. and R.M. conceived and designed the study, developed methodologies, and wrote the manuscript. C.S. performed most of the experiments and data analyses. K.M. developed and provided methodologies and technical advice. K.K. performed part of the data analyses. L.Y. performed part of the experiments. R.-S.N. provided methodologies and technical advice. R.M. supervised the study and secured funding. All authors discussed the results and contributed to manuscript editing.

## Extended Data

**Extended Data Fig. 1.**
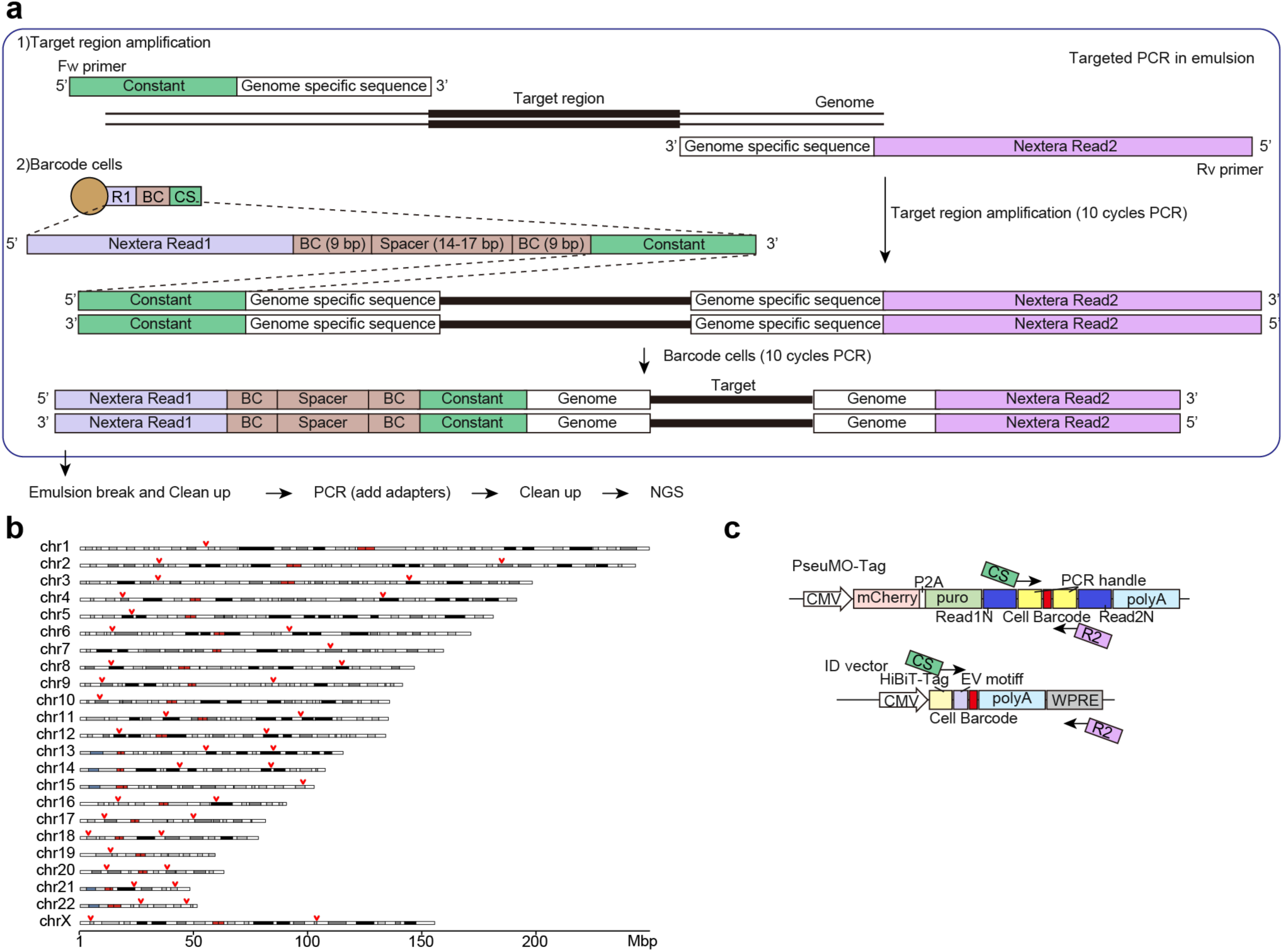
Workflow for quantifying ecDNA copy-number distribution by scDNA-seq. a, Schematic of the scDNA-seq experimental workflow. The target regions were amplified, followed by barcode assignment within the emulsions. The emulsions were broken, adapters were added, and libraries were prepared for NGS. **b,** Reference regions used for normalization are indicated by red arrows. One or two reference regions were designed for each chromosome. **c,** Schematic of the two types of celularl barcodes, PseuMO-Tag and ID vector.

**Extended Data Fig. 2.**
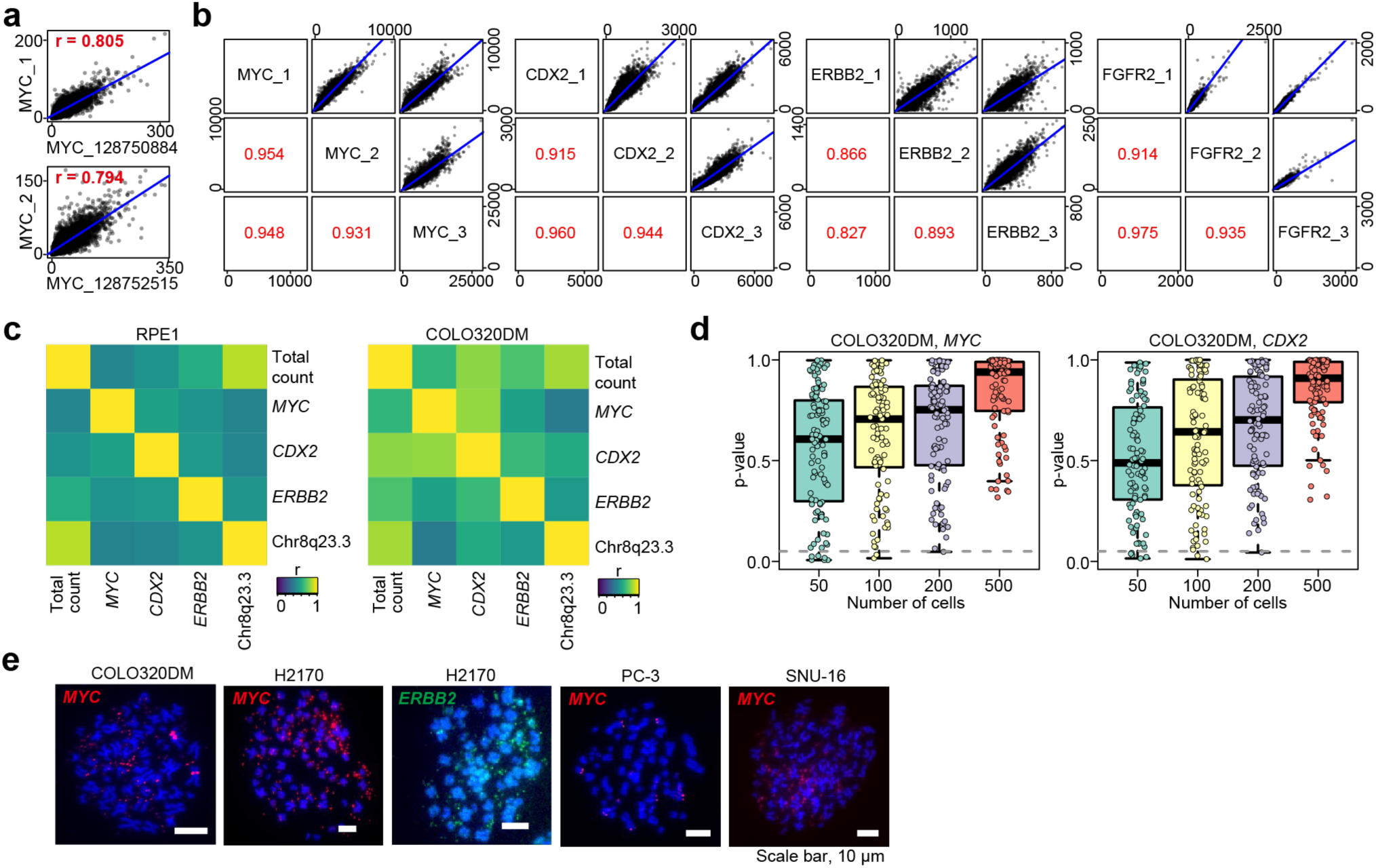
Quality check of the ecDNA copy-number distribution quantification framework. a, Correlation of the read counts between amplicons generated with the commercially available Myeloid Panel v2 primers and those generated with custom primers targeting the same regions. **b,** Correlation of the read counts between amplicons generated with primers designed to amplify different regions of the same gene. **c,** Correlation of the read counts among genes in RPE1 (left) and COLO320DM (right). In COLO320DM, *MYC* and *CDX2* are located on the ecDNA. **d,** Box plot showing the distribution of p-values from 100 iterations of the KS test. For each iteration, the indicated numbers of cells were randomly sampled from 1,266 COLO320DM cells to test whether the distributions of *MYC* and *CDX2* copy numbers differed from those of the original population. **e,** Representative FISH images of each cell line. *MYC* and *ERBB2* are shown in red and green, respectively.

**Extended Data Fig. 3.**
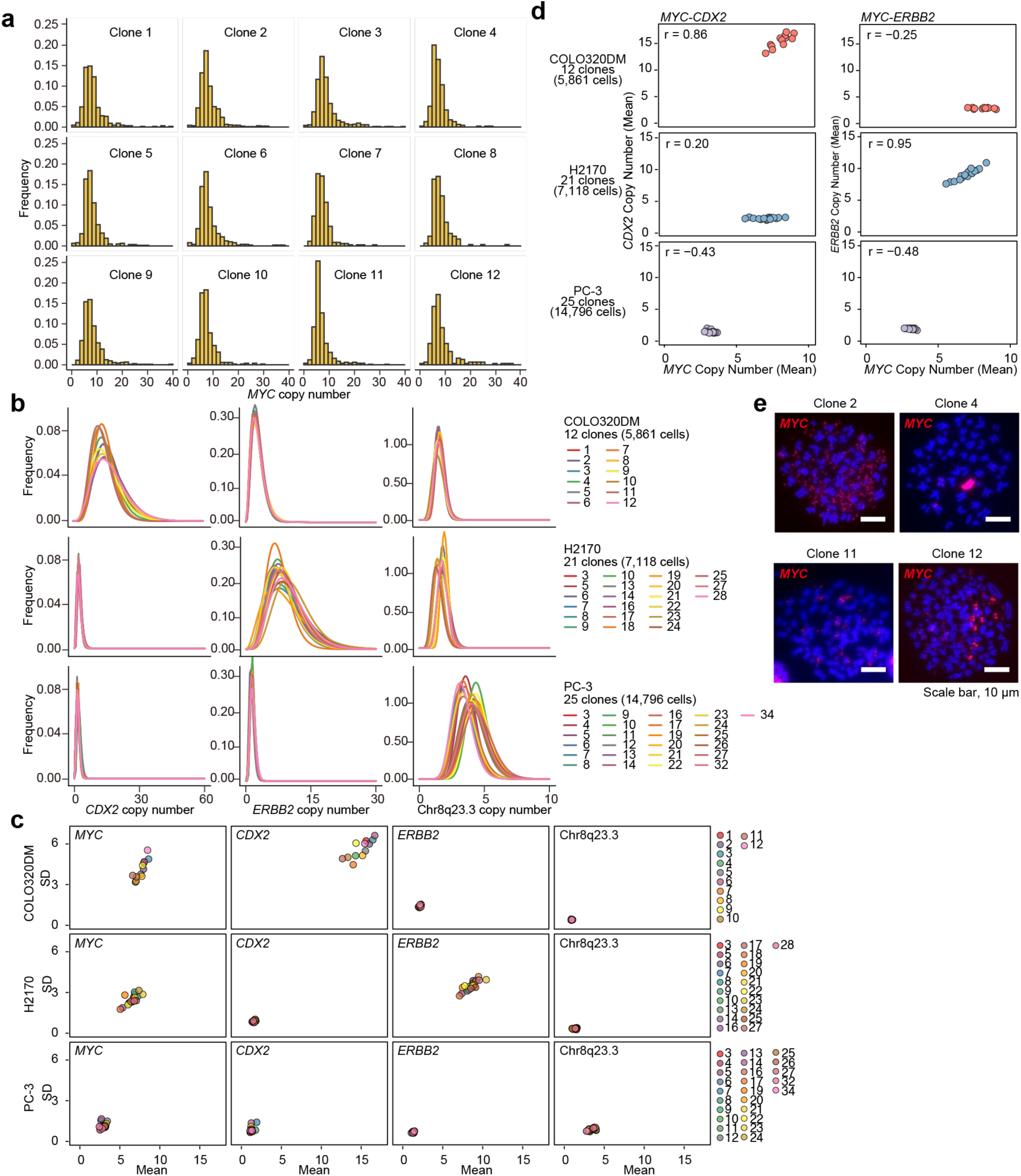
Copy-number distributions of clones from three cell lines. a, Histograms of copy-number distributions of *MYC* in 12 clones of COLO320DM. **b,** Gamma distribution fitting of copy-number distributions of *CDX2*, *ERBB2*, and chr8q23.3 for each clone derived from COLO320DM, H2170, and PC-3 cells. **c,** Scatter plot of the mean and SD values for *MYC*, *CDX2*, *ERBB2*, and chr8q23.3 for each clone derived from COLO320DM, H2170, and PC-3 cells. **d,** Correlation between the mean *MYC* copy number and the mean *ERBB2* copy number for each clone derived from COLO320DM, H2170, and PC-3 cells. **e,** Representative FISH images of COLO320DM clone 2, 4, 11, and 12.

**Extended Data Fig. 4.**
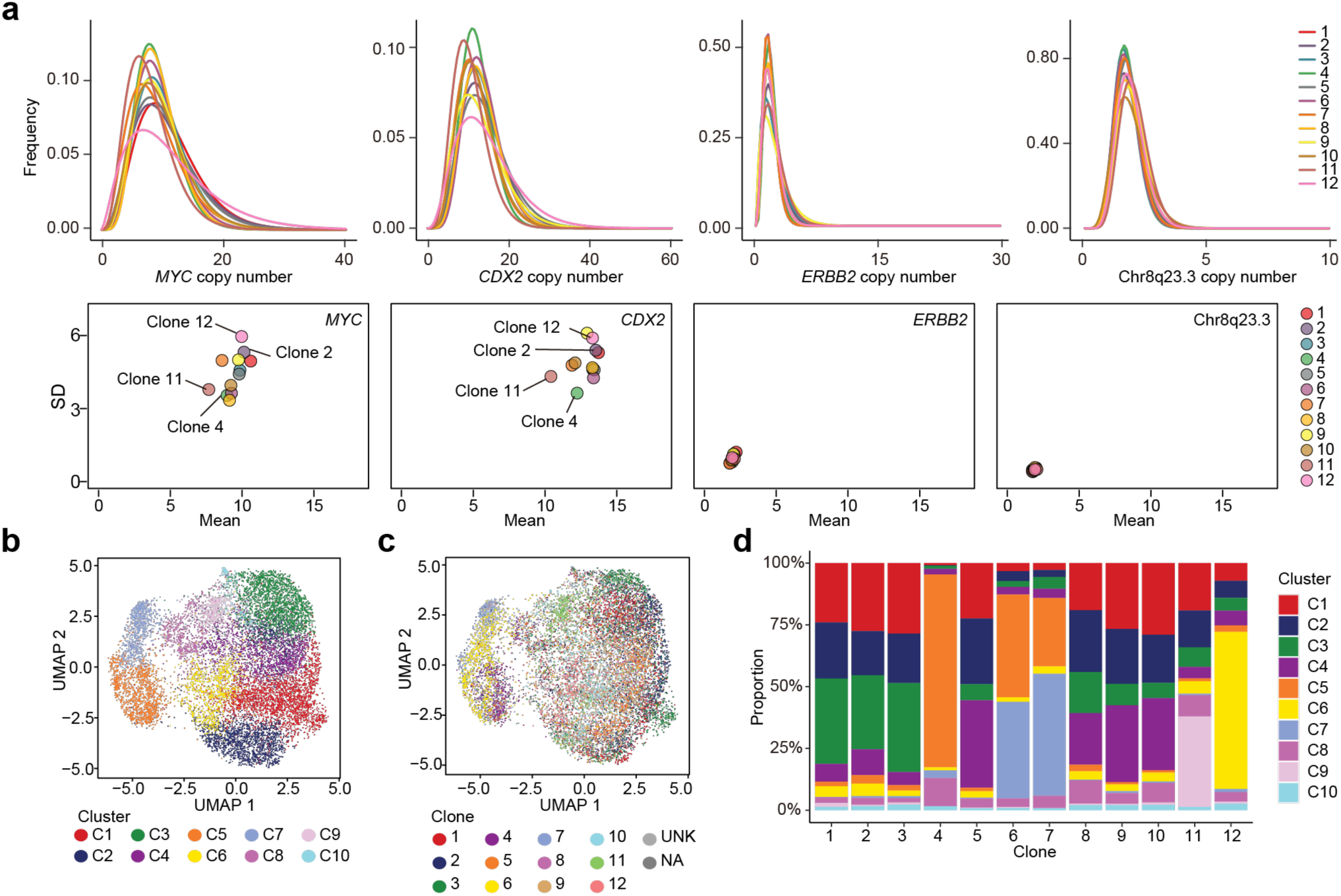
Clonal diversity of each COLO320DM clone. a, Copy-number distributions and scatter plot of the mean and SD values for *MYC*, *CDX2*, *ERBB2*, and chr8q23.3 in each COLO320DM clone after 18 additional cell divisions, from the time point of the data shown in Fig. 2b– d and Extended Data Fig. 3. The dataset includes 7,220 cells. **b,** UMAP representation of 12 COLO320DM clones, colored by clusters. **c,** UMAP representation of 12 COLO320DM clones before additional cell divisions, colored by clones. **d,** Stacked bar plot showing the proportion of each cluster assigned to each clone.

**Extended Data Fig. 5.**
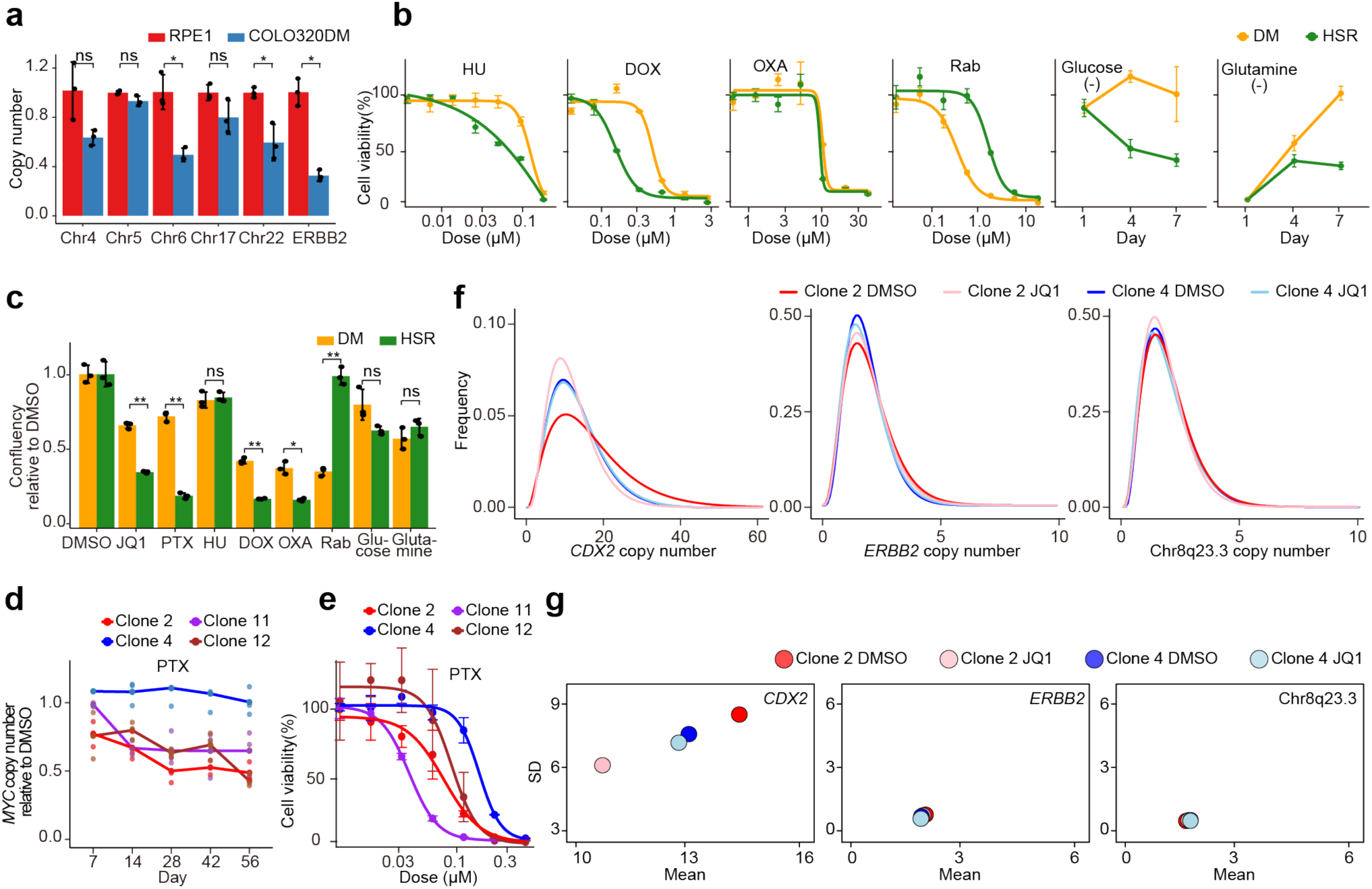
Clone-specific reduction of ecDNA copy number under JQ1 and PTX. a, Copy number of the indicated regions in COLO320DM cells, measured by quantitative PCR (qPCR) and normalized to that in RPE1 cells. Data are mean ± SD (n = 3). RPE1 is shown in red and COLO320DM in blue. *p* = 0.095 (Chr4), *p* = 0.090 (Chr5q23.1), \**p* = 0.015 (Chr6), *p* = 0.11 (Chr17), \**p* = 0.033 (Chr22), and \**p* = 0.0038 (*ERBB2*). **b,** Cell viability assay of COLO320DM and COLO320HSR under the same conditions shown in Fig. 3a. Data are mean ± SD (n = 3). **c,** \*\**p* < 0.001 (JQ1), \*\**p* < 0.001 (PTX), *p* = 0.65 (HU, hydroxyurea), \*\**p* = 0.0016 (DOX, doxorubicin), \**p* = 0.011 (OXA, oxaliplatin), **p < 0.0014 (Rab, rabusertib), *p* = 0.093 (Glucose; glucose depletion), and *p* = 0.21 (Glutamine; glutamine depletion). **d,** Time-dependent change in *MYC* copy number for COLO320DM clones 2, 4, 11, and 12 treated with 100 nM PTX over time, measured by qPCR, and normalized to the corresponding DMSO-treated control for each clone. Each point represents an individual measurement, and the lines indicate the mean values (n = 3). **e,** Cell viability assay of COLO320DM clones 2, 4, 11, and 12 treated with PTX. Data are mean ± SD (n = 3). **f-g,** Gamma distribution fitting (**f**), and scatter plot of the mean and SD values (**g**) for the copy number of *CDX2*, *ERBB2*, and chr8q23.3 in COLO320DM clone 2 and 4 after two-weeks of DMSO or JQ1 treatment.

**Extended Data Fig. 6.**
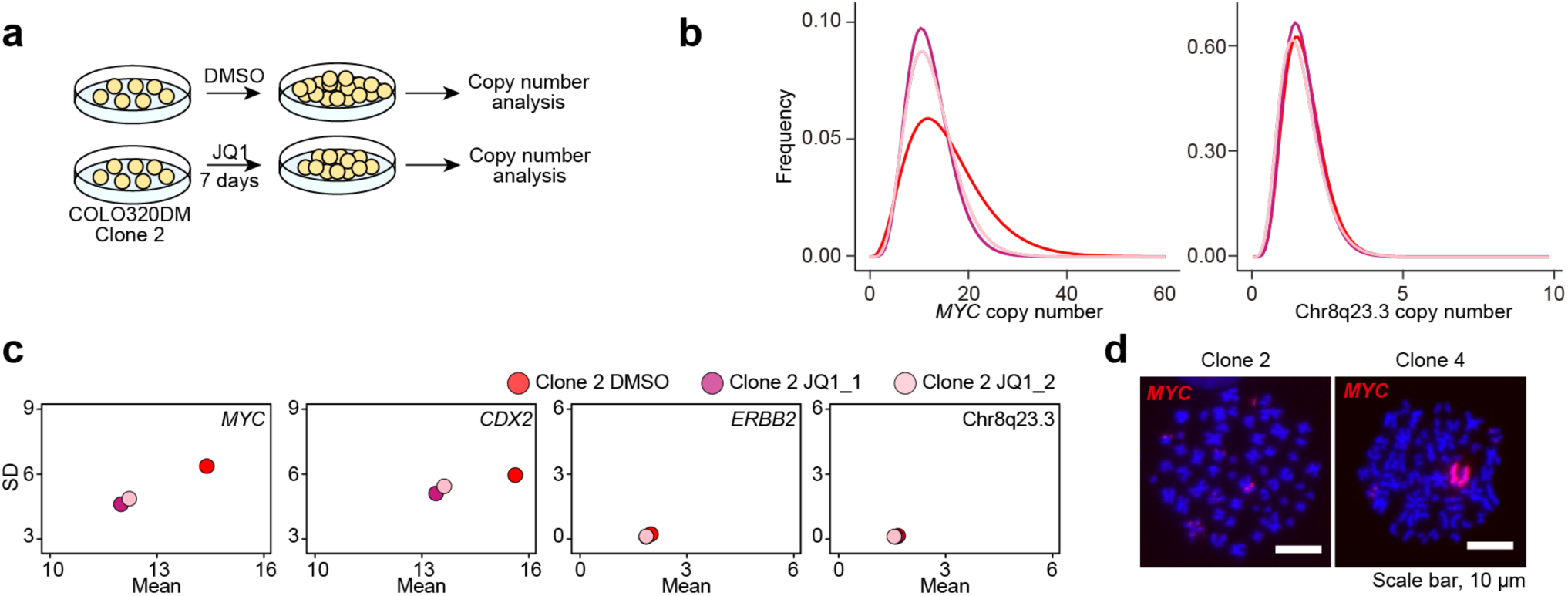
Clone-specific reduction of ecDNA copy number following short-term JQ1 treatment. a, Schematic of the experimental design for examining ecDNA copy-number distributions changes after JQ1 treatment using scDNA-seq analysis in COLO320DM clones 2. The duration of JQ1 treatment was one week. **b,** Gamma distributions fitting of copy-number distributions of *MYC* and chr8q23.3 in COLO320DM clones 2 after one-week DMSO or JQ1 treatment. **c,** Scatter plot of the mean and SD values of copy-number for *MYC*, *CDX2*, *ERBB2*, and chr8q23.3 in COLO320DM clone 2 after one-week DMSO or JQ1 treatment. **d,** Representative FISH images of COLO320DM clones 2 and 4 after 56 days of JQ1 treatment.

**Extended Data Fig. 7.**
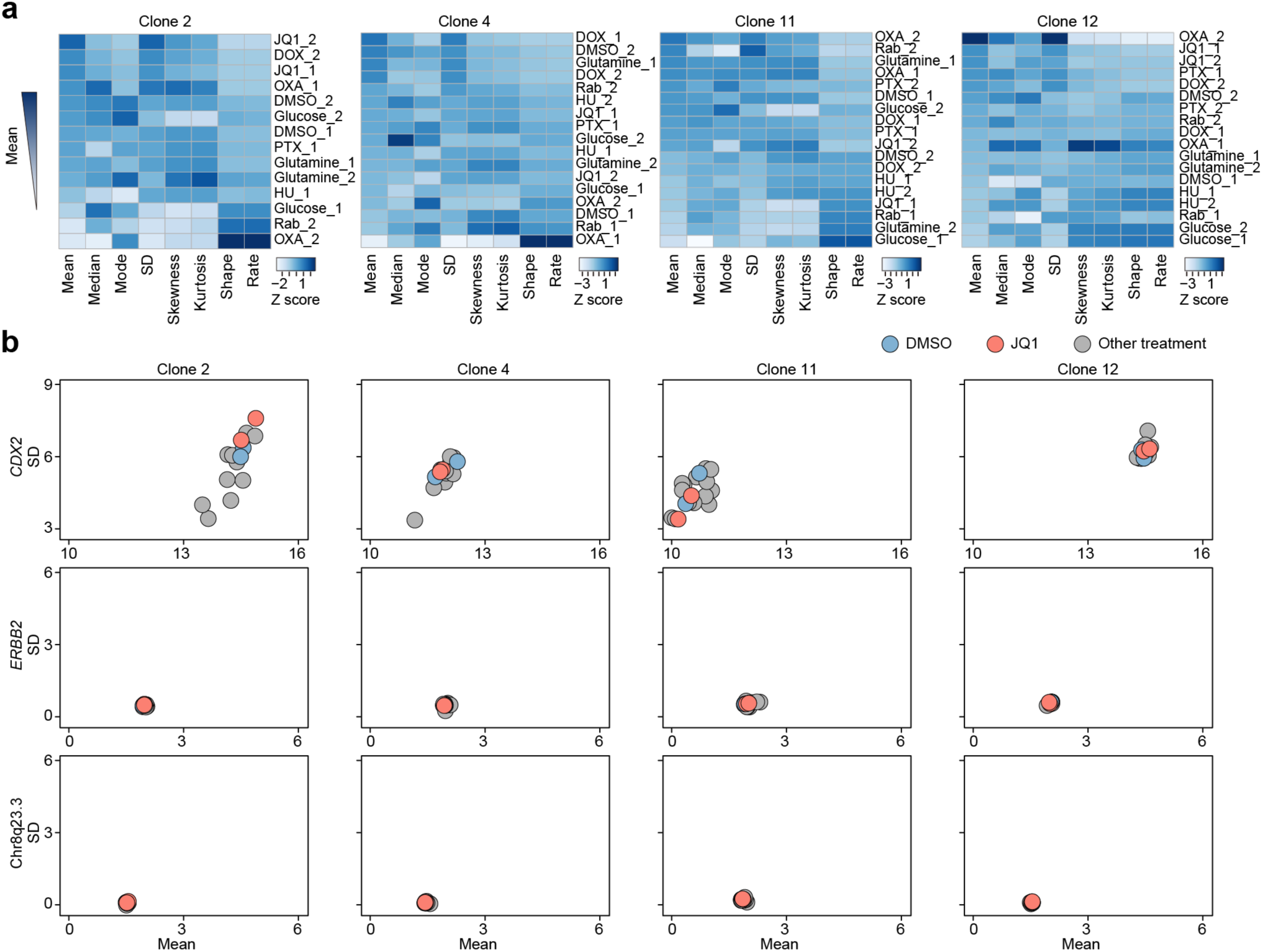
Clonal diversity of ecDNA copy number changes in drug responses. a, Heatmap of eight parameters of *MYC* in each clone, sorted in descending order of the mean value. Each column was normalized by Z-score scaling. b, Scatter plot of the mean and SD values of copy number for *MYC*, *CDX2*, *ERBB2*, and chr8q23.3 in COLO320DM clones. The DMSO-treated group is shown in blue and the JQ1-treated group in red.

**Extended Data Figure Fig. 8.**
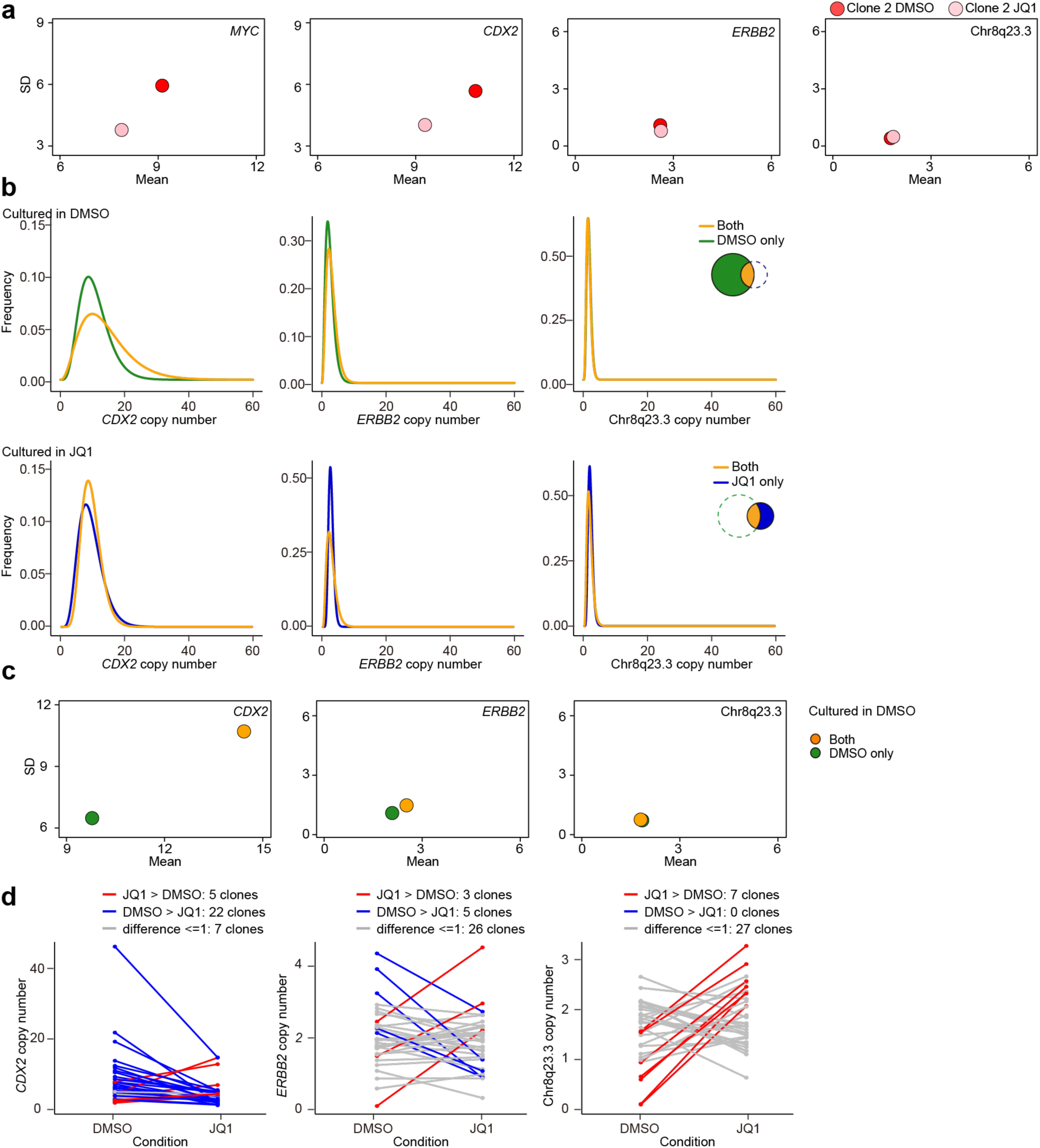
Active reconfiguration of ecDNA in clone 2 after JQ1 treatment. a, Gamma distribution fitting of copy-number distributions of *MYC*, *CDX2*, *ERBB2*, and chr8q23.3 in COLO320DM clone 2 after 10 days DMSO treatment (red) and 24 days JQ1 treatment (pink). **b,** Gamma distribution fitting of copy-number distributions of *CDX2*, *ERBB2*, and chr8q23.3 copy-number distributions under DMSO (top) and JQ1 (bottom) conditions. Copy-number distributions of clones detected only in the DMSO-treated group are shown in green (top), those detected only in the JQ1-treated group are shown in blue (bottom), and clones detected in both groups are shown in orange. **c,** Scatter plot of the mean and SD values for *CDX2*, *ERBB2*, and chr8q23.3 copy number in the DMSO-treated group. Clones detected only in the DMSO-treated group are shown in green, and clones detected in both groups are shown in orange. **d,** Comparison of the mean *CDX2*, *ERBB2*, and chr8q23.3 copy number between the DMSO and JQ1 culture conditions in clones detected in both the DMSO- and JQ1-treated groups.

**Extended Data Figure Fig. 9.**
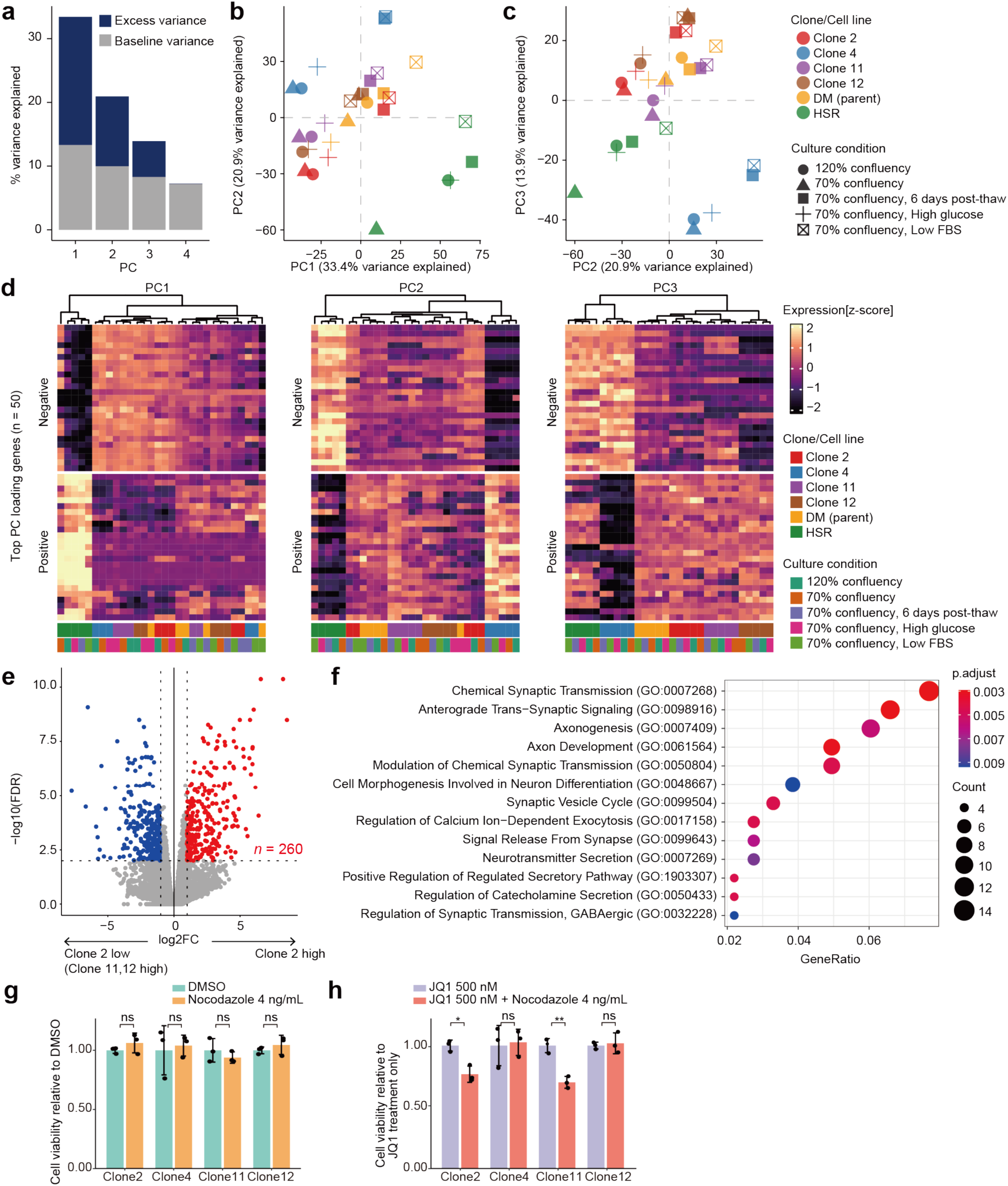
Clone 2 exhibits neuron-like transcriptional characteristics. a, Bar plot showing the percentage of variance explained by the first to fourth principal components (PCs). The observed values from PC1 to PC3 exceed the baseline values predicted by the broken-stick model. **b–c,** Principal component (PC) plots of the gene expression profiles of COLO320DM clones 2, 4, 11, 12, the parental line, and COLO320HSR under various culture conditions. PC1 vs. PC2 is shown in (b), and PC2 vs. PC3 in (c). **d,** Heatmap showing the expression of the top 25 positively and 25 negatively loaded genes for PC1, PC2, and PC3. **e,** Volcano plot showing the differential gene expression between clone 2 and clones 11 and 12. Each dot represents a gene. Red dots indicate genes significantly upregulated in clone 2, while blue dots indicate genes significantly upregulated in clones 11 and 12. **f,** Dot plot showing the gene ontology analysis of genes highly expressed in clone 2. **g,** Cell viability assays of each clone after 7-day treatment with DMSO or 4 ng/mL nocodazole. Data are mean ± SD (n = 3). *p* = 0.33 (clone2), *p* = 0.78 (clone4), *p* = 0.41 (clone11), and *p* = 0.44 (clone12). **h,** Cell viability assays of each clone after 7-day treatment with 500 nM JQ1 or 500 nM JQ1 plus 4 ng/mL nocodazole. Data are mean ± SD (n = 3). \**p* = 0.010 (clone2), *p* = 0.83 (clone4), * \**p* = 0.0026 (clone11), and *p* = 0.75 (clone12).

## Supplementary Information

**Supplementary Note. 1** Establishing a quantitative basis for the ecDNA copy-number distribution. **Supplementary Note. 2** Optimizing the description and parameterization of copy-number distribution.

**Supplementary Table 1.** Primers used in the experiments.

**Supplementary Table 2.** Primers used as for validation of primer efficiency.

**Supplementary Table 3.** Goodness of fit for each distribution evaluated by Akaike Information Criterion

**Supplementary Table 4.** Differential gene expression analysis comparing clone 2 with clone 11 and 12

