## Supplementary Note 1., Supplementary Note 2. for "Plasticity of extrachromosomal DNA segregation during drug adaptation"

**Supplementary Note 1. Establishing a quantitative basis for ecDNA copy-number distribution**

Regarding quantitative framework established here for ecDNA copy-number analysis, targeted PCR was performed on emulsion droplets followed by sequencing (**Extended Data Figure 1a**). Read counts were normalized using the diploid cell line RPE1 as a reference. Control regions (n = 40) were designed in the intergenic regions and distributed evenly across the chromosomes (**Extended Data Figure 1b**). Normalization was done using according to the equation described below and detailed in the Methods section.

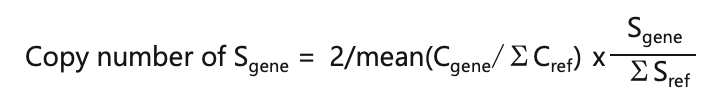

C_gene_; Read counts of the target region in RPE1 cells

C_ref_; Read counts of the reference region in RPE1 cells

S_gene_; Read counts of the target region in ecDNA-positive cells

S_ref_; Read counts of the reference region in ecDNA-positive cells

All targeted PCR primer sets were custom-designed. This strategy enables the dynamic adjustment of the target regions based on the experimental requirements and reduces sequencing costs by restricting amplification to selected loci. Details of the primer design are provided in the Methods section. To determine whether custom-designed primers could amplify target regions with an efficiency comparable to commercially available primer panels, custom primers and the corresponding panel primers were mixed and run together in the same reaction. The x-axis represents the read counts obtained using the commercially available Myeloid Panel v2 primer panel, and the y-axis represents those obtained by the custom-designed primers. The read counts exhibited a strong correlation, with a Pearson’s r value of approximately 0.8, indicating comparable amplification efficiency between the two primer sets (**Extended Data Figure 2a**). Read counts for the amplicons generated with primers designed for different regions within the same gene exhibited strong correlations, indicating consistent amplification efficiency among the primer sets targeting the same gene (**Extended Data Figure 2b**). Moreover, in COLO320DM cells, copy numbers of *MYC* and *CDX2*, both located on ecDNA, were moderately correlated, whereas no correlation was evident in RPE1 cells (**Extended Data Figure 2c**).

**Supplementary Note 2. Optimizing the description and parameterization for copy-number distribution**

To establish the optimal method to describe and categorize the differences among the samples, we addressed the following three questions: (i) how many cells are required to accurately reconstruct the distribution; (ii) which statistical distribution best approximates the copy-number histograms; and (iii) which metrics are most appropriate for quantitatively evaluating the distribution shapes.

To address question (i), we conducted the following analysis using the *MYC* and *CDX2* copy-number data from 1,266 COLO320DM cells (Figure 1b). We randomly sampled 50 cells **without replacement** and compared their distributions with that of the entire population using the Kolmogorov–Smirnov test. Sampling and comparison were repeated 100 times, and the resulting p-values were calculated. The same procedure was conducted for samples containing 100, 200, and 500 cells. The distributions of the p-values are shown as box plots (**Extended Data Figure 2d**). As the number of sampled cells increased, the distributions more closely resembled that of the population. For both *MYC* and *CDX2*, sampling 50 cells yielded distributions that were already largely similar to that of the parental population, suggesting that this sample size was acceptable. However, sampling 100 cells provided even more consistent results, with only two out of 100 tests yielding p-values below 0.05. Therefore, we concluded that data from at least 100 cells are preferable to accurately capture the true distribution.

To address question (ii), we determined which statistical distribution best approximates the observed copy-number distributions. Because previous studies have indicated that ecDNAs are randomly segregated, the overall distributions were expected to be symmetric; however, the histograms shown in Figure 1b showed right-skewed tails. Therefore, we fitted the copy-number data for *MYC* and *CDX2* in COLO320DM, *MYC* and *ERBB2* in H2170, *MYC* in SNU-16, and *MYC* in PC-3 to normal, gamma, and log-normal distributions. The goodness of fit was evaluated using the Akaike Information Criterion (AIC). The goodness of fit for the gamma and log-normal distributions was comparable (**Supplementary Table 3**). Therefore, we adopted the gamma distribution for subsequent analyses, because its parameters (shape and rate) facilitate direct comparison and visualization, and the copy-number variation arises from stochastic partitioning rather than multiplicative amplification (**Figure 2c**).

To address question (iii), we considered several quantitative indicators to describe the distribution shape: mean, median, and mode (representing copy-number level); standard deviation (SD; representing dispersion); skewness (asymmetry); kurtosis (peakedness); and the gamma distribution parameters, shape and rate. As shown in Figure 1e, the mean, median, and mode were highly correlated. Moreover, the skewness, kurtosis, and the gamma parameters, shape and rate, largely depend on the values of the mean and SD because these parameters are mathematically derived from, or closely related to, the mean and SD. Therefore, we concluded that the mean and SD are the most informative and concise metrics for evaluating differences in distribution shape.
